# MEG cortical microstates: spatiotemporal characteristics, dynamic functional connectivity and stimulus-evoked responses

**DOI:** 10.1101/2021.03.25.436979

**Authors:** Luke Tait, Jiaxiang Zhang

## Abstract

EEG microstate analysis is a useful approach for studying brain states - nicknamed ‘atoms of thought’ - and their fast transitions in healthy cognition and disease. A key limitation of conventional microstate analysis is that it must be performed at the sensor level, and therefore gives limited anatomical insight into the cortical mechanisms underpinning these states. In this study, we generalise the microstate methodology to be applicable to source-reconstructed electrophysiological data. Using simulations of a neural-mass network model, we first established the validity and robustness of the proposed method. Using MEG resting-state data, we uncovered ten microstates with distinct spatial distributions of cortical activation. Multivariate pattern analysis demonstrated that source-level MEG microstates were associated with distinct functional connectivity patterns. Using a passive auditory paradigm, we further demonstrated that the occurrence probability of MEG microstates were altered by evoked auditory responses, exhibiting a hyperactivity of the microstate including the auditory cortex. Our results support the use of MEG source-level microstates as a data-driven method for investigating brain dynamic activity and connectivity at the millisecond scale.

## 1 Introduction

Whole-brain, non-invasive functional neuroimaging of the human brain is a useful tool for uncovering the mechanisms underpinning cognitive functions and neurological disease (van den Heuvel and Hulshoff Pol, 2010; Douw et al., 2011; Babiloni et al., 2016; Cohen, 2018; Michel and Koenig, 2018; Lee et al., 2019). In recent years, there has been much evidence that even at rest the activity of the human brain is highly dynamic, transitioning between a small number of functional brain-states with specific patterns of activation or synchrony across the cortex (Smith et al., 2009; Baker et al., 2014; Khanna et al., 2015; Michel and Koenig, 2018; O’Neill et al., 2018; Taghia et al., 2018; Vidaurre et al., 2018). These functional brain states have been associated with a range of cognitive domains and levels of consciousness (Smith et al., 2009; Brodbeck et al., 2012; Britz et al., 2014; Milz et al., 2016; Seitzman et al., 2017; Liégeois et al., 2019; Zappasodi et al., 2019; Zhou et al., 2019), demonstrating that more advanced insight into the non-stationarity of brain stats and their dynamics may be crucial to understanding cognition and disease.

In functional MRI (fMRI) data, techniques to study functional brain-states have included independent component analysis (ICA) (Beckmann et al., 2005; Smith et al., 2009) and sliding-window analysis (Allen et al., 2014), uncovering a small number of reproducible networks known as resting-state networks (RSNs) (Smith et al., 2009). Temporal resolution of fMRI is limited by the slow dynamics of the haemodynamic response function, meaning transitions between states on a sub-second scale are likely to be missed. ICA has also been used in EEG and MEG to recreate fMRI-RSNs (Brookes et al., 2011; Liu et al., 2017), but to do so this is typically performed on a slower time scale matching those of fMRI through analysis of downsampled power envelopes, losing the high temporal resolution and oscillatory phase information gained from the use of EEG/MEG. It follows that alternative methods are required to uncover faster time-scale (on the order of milliseconds) dynamics in EEG/MEG data (O’Neill et al., 2018), and it is currently unclear the extent to which spatiotemporal properties of millisecond scale non-stationarity reflect those of slower RSNs.

Other methods for EEG/MEG analyses are available to examine to rapid changes in brain-state. One is to apply sliding window analysis to the EEG/MEG time courses (de Pasquale et al., 2010; Brookes et al., 2014; O’Neill et al., 2015; de Pasquale et al., 2016; Lopes et al., 2020) and subsequently cluster functional networks across windows (Allen et al., 2014; O’Neill et al., 2015; Mheich et al., 2015; Hassan et al., 2015). This approach has the limitation of the need for an arbitrary *a priori* selected window size: too short windows lead to results susceptible to noise, while too large windows result in non-stationarity at fast time scales being missed (O’Neill et al., 2018). Indeed, there is no consensus on optimal sliding window length and the literature has covered a wide range, from 25ms to 30s (O’Neill et al., 2018). A number of alternative approaches to sliding windows exist (O’Neill et al., 2018), including Hidden Markov Models (HMMs) (Baker et al., 2014; Vidaurre et al., 2018). HMMs rely on assumptions about the underlying process of state transitions, both in a detailed generative model and Markovian transitioning between states. It is unclear whether these assumptions are always met by spontaneous transitions between brain states (Gschwind et al., 2015).

Another alternative approach to study dynamics of functional brain-states, the focus of the current study, is microstate analysis (Khanna et al., 2014; Michel and Koenig, 2018). Conventional EEG microstate analysis involves clustering the sensor-space spatial topographies using algorithms such as *k*-means or hierarchical clustering (Khanna et al., 2015; von Wegner et al., 2018). Microstate analysis therefore does not require an arbitrarily chosen window length, and has minimal assumptions about the underlying generative process. Resting-state EEG microstates are robust and highly reproducible (Michel and Koenig, 2018), and have been associated with fMRI resting-state networks (Britz et al., 2010; Musso et al., 2010; Yuan et al., 2012; Schumacher et al., 2019; Abreu et al., 2020; Xu et al., 2020; Zoubi et al., 2020) and cognitive domains (Brodbeck et al., 2012; Britz et al., 2014; Milz et al., 2016; Seitzman et al., 2017; Zappasodi et al., 2019), earning EEG microstates the nickname the ‘atoms of thought’ (Lehmann, 1990). EEG microstates have also been demonstrated to be a potentially useful clinical tool for understanding and diagnosing neurological diseases such as Alzheimer’s disease and other dementias (Nishida et al., 2013; Musaeus et al., 2019; Smailovic et al., 2019; Schumacher et al., 2019; Tait et al., 2020b), schizophrenia (Lehmann et al., 2005; Andreou et al., 2014; Tomescu et al., 2014), and a range of other disorders (Khanna et al., 2014).

However, a key limitation of sensor-space microstate analysis is anatomical interpretation. A number studies have attempted to reconstruct the electrophysiological sources underpinning microstates maps by combining sensor-space microstate analysis with subsequent source reconstruction (Custo et al., 2014, 2017; Pascual-Marqui et al., 2014; Milz et al., 2016, 2017; Tait et al., 2020b). However, this approach may be limiting insight into functional brain states for a number of reasons. Firstly, since the inverse problem is not unique, it is possible that different spatial patterns of brain activation may give rise to similar topographical maps, and therefore each sensor-level microstate may be associated with more than one active network. Secondly, due to the spatial blurring in EEG as a result of volume conduction through tissues of different conductivities, EEG microstate topographies have low spatial resolution and potentially cannot differentiate between finer differences between maps; indeed, EEG microstate topographies are reliable with as few as eight electrodes (Khanna et al., 2014), suggesting the topographies do not contain much spatial detail. Finally, since alpha band occipital sources dominate the sensor-space eyes-closed resting-state EEG (Kropotov, 2009) likely due to head shape and the forward model resulting in high signal-to-noise ratio for these regions (Goldenholz et al., 2009), it is likely that these same sources predominantly determine the microstate topographies (Milz et al., 2017), therefore suggesting the sensor-space EEG microstate maps may be under-weighting the importance of non-occipital or non-alpha-band networks.

Instead of performing microstate analysis in sensor space and then subsequently projecting to source space, here we propose first projecting M/EEG data to source space and subsequently clustering the source dynamics. Unfortunately, some steps of the the EEG microstate pipeline means it is not directly applicable to source-reconstructed recordings, such as relying the requirement to re-reference to average and being unable to account (at the group level) for arbitrary source flipping. Here we adapt methodology and study microstates in MEG source space at rest and during task. We hypothesise that each microstate is associated with distinct patterns of functional connectivity across the cortex and use machine learning to test this hypothesis. Finally, we demonstrate that source-space microstate features are dependent of cognitive state and examine the microstate response to auditory stimuli.

## 2 Methods

### 2.1 A source-space microstate pipeline

The source-space microstate segmentation pipeline used here is based upon the widely used EEG sensor-space *k*-means pipeline presented by Pascual-Marqui et al. (1995) and Koenig et al. (1999), and is outlined in Table 1.

**Table 1:** The microstate pipeline, generalized for different recording modalities. Details and justification of these steps are given in section 2.1.

| **Algorithm: Source-space microstate pipeline** |  |
| --- | --- |
| 1 | Source reconstruct sensor data, band-pass filter, and parcellate |
| 2 | Extract activity patterns (maps) at GFP peaks |
| 3 | Run $k$-means clustering on transformed (absolute value) maps. |
| 4 | Backfit cluster centroids to data |
| 5 | Calculate microstate statistics. |

However, there are a number of steps in the sensor-EEG microstate pipeline which limit application to source-space data. One such limitation is the requirement to re-reference to average; while the EEG data itself does not necessarily have to be referenced to average, the pipeline relies on metrics such as standard deviation of the signal across sensors (global-field power, GFP) and correlation between spatial topographies, both of which involve subtracting the mean of the map (i.e. common average re-referencing). The pipeline presented here is modified, replacing standard deviations and correlations with (normalized) vector norms and cosine similarity respectively. These metrics are equivalent in the case of zero mean data, and hence the methodology presented here is a generalization of the sensor-EEG pipeline which may be used on reference-free data such as source data or sensor MEG, while exactly preserving the EEG-microstate pipeline is under the requirement that data is first re-referenced to common average.

A second issue for source reconstructed data is that of source flipping. At the voxel level, any dipole can arbitrarily be ‘flipped’ by changing the sign of both the dipole orientation and time course. While in this study we work on the level of ROIs as opposed to individual dipoles, if parcellation is performed via a single voxel with maximal variance dipole flipping is still an issue. Another common method of parcellation (used in this study) is to take the first principal component of all dipoles within an ROI, in which case the sign of both the time course and spatial topography of the first principal component may be flipped to obtain an identical result. This source flipping is not problematic for analysis of a single M/EEG scan, but for group level analyses different participants may have different spatial patterns of source flipping. For *N* ROIs, source flipping can result in 2^*N−*1^ possible combinations. For 230 ROIs, used in this study, source flipping could result in a number of possible topographies of the order 10^68^ for a single microstate. Therefore applying traditional EEG microstate analysis to source signals may result in an overestimate of the true number of clusters in group level analysis. To confound this issue, one may use either the amplitude envelope or take the (element-wise) absolute value of the samples. The former requires a narrow band signal and rejects phase information, while the latter can be applied to broadband data and mostly maintains phase information (excluding phase differences of *π*). Recent work has highlighted that phase information is likely crucial for encoding microstate sequences (von Wegner et al., 2021). For these reasons, and based on the results of simulations (section 3.1), throughout this manuscript we use the absolute value of source estimates during clustering and map fitting.

A description of the source microstate pipeline are given in the following sections. While these sections give an overview the source-space pipeline, mathematical details and a generalized description of the pipeline for different modalities based appropriate modality-specific spatial transformation of the data (e.g. re-reference to average for sensor-EEG, in which case the traditional sensor-EEG microstate pipeline is recovered exactly) are given in Supplementary Text S1.1.

#### 2.1.1 Source-reconstruction and parcellation

The aim of this manuscript is to present a pipeline for source-spaced M/EEG microstate analysis. The first step of the pipeline is therefore source reconstruction of the M/EEG data. A range of methodologies for source reconstruction are currently available and the choice of methodology chosen should be data specific (Tait et al., 2020a). Here, eLORETA (Pascual-Marqui, 2007, 2009) was used for source-reconstruction. Full details of the pipeline for source-reconstruction in this study are given in section 2.5.3. Since distributed source reconstruction methodologies typically reconstruct a large number of dipoles (several thousand), a key step following source reconstruction is spatial downsampling to increase tractability for spatial clustering. This was done through parcellation of source time courses into regions of interest. Here, the HCP230 atlas (Tait et al., 2020a), a version of the Human Connectome Project’s multimodal parcellation Glasser et al. (2016) optimized for resting-state MEG, was chosen to reduce the source data from approximately 10,000 voxels to 230 ROI time courses. Source-reconstructed data was band-pass filtered 1-30 Hz in line with sensor-EEG microstate studies (Michel and Koenig, 2018).

#### 2.1.2 Extraction of GFP peaks

Subsequently, samples with optimal signal-to-noise ratio (SNR) and topographic stability were chosen for clustering. These samples correspond to those with peaks in the global field power (GFP) (Supplementary Figure S4; Michel et al. (2009), Koenig and Brandeis (2016)). In sensor-EEG microstate analysis, the GFP at a time sample *t* is defined as the standard deviation of the electric potential across sensors. For source-reconstructed data, the GFP is given by the vector norm of the signal across ROIs and hence is the total deviation from zero current source density (which is equivalent to the standard deviation in the case of zero mean).

Due to the 1-30 Hz bandpass fitler, the GFP time course is temporally smooth, and hence local maxima of the GFP (i.e. GFP peaks) can be found using a peak-finding algorithm. Here, Matlab’s findpeaks function was used to identify GFP peaks.

For group level analysis in this study, we randomly sampled and concatenated 5000 GFP peaks per participant. For each condition (rest/task), two MEG scans were available for each participant (*n* = 30) recorded on different dates (see section 2.5), and here we only extracted the 5000 GFP peaks from the first scan (allowing the second scan to be used for validation purposes). Hence, across participants we extracted 150,000 GFP peaks. Spatial maps at each GFP peak were submitted for *k*-means clustering.

#### 2.1.3 *k*-means clustering

The next step of the pipeline is to cluster the submitted maps. For a given number of states, *k*, the algorithm for *k*-means clustering is outlined in Table 2, while a clear graphical representation is given in Koenig et al. (1999) and Michel et al. (2009).

**Table 2:** Outline of the *k*-means algorithm.

| **Algorithm: $k$-means clustering** |  |
| --- | --- |
| 1 | Select $k$ maps as initial centroids |
| *While not converged:* |  |
| 2 | Calculate distance between maps and centroids |
| 3 | Cluster each map based on nearest centroid |
| 4 | Calculate new centroids from all maps in cluster |
| 5 | Test for convergence (no change in clusters) |
| *end while* |  |

Firstly, *k* maps are chosen as the initial cluster centroids. Here, the *k*-means++ algorithm was used for selection of initial maps (Arthur and Vassilvitskii, 2007). In the *k*-means algorithm, the distance between each submitted map and each centroid is calculated using the cosine distance. In the case of zero mean (e.g. such as average re-referenced EEG instead of source data), the cosine distance is equal to one minus correlation, which is the metric used in the sensor-EEG *k*-means algorithm. Each map is subsequently labelled as belonging to the state with the closest centroid. In sensor-EEG, new cluster centroids are calculated as the first principal component of all maps within the cluster (Pascual-Marqui et al., 1995). In source-space, new cluster centroids are calculated as the eigenvector corresponding to the largest eigenvalue of the matrix 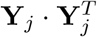, where rows of the matrix **Y**_*j*_ are the maps within cluster *j*. This eigenvector is equal to the first principal component in the case of zero mean. Using the new centroids, the procedure of calculating cluster labels and updating cluster centroids is iterated until convergence is reached. Due to random initial seeding, the *k*-means algorithm was repeated 20 times and the repetition with highest global explained variance (GEV) (Murray et al., 2008) was chosen for further analysis (Koenig et al., 1999).

The choice of number of states *k* is a free parameter. In this study, we used the kneedle algorithm (Satopä ä et al., 2011) to determine the optimum number of states. As the number of clusters *k* increases, so does the GEV. However, increasing *k* above the true number of states will only give marginal increases to GEV, and therefore the plot of GEV vs *k* has a characteristic ‘knee’ shape. The kneedle algorithm aims to find the number of clusters in the dataset by finding the value of *k* for which this knee occurs.

#### 2.1.4 Backfitting to source time courses

After identifying the microstate maps from the sample of 150,000 GFP peaks, each sample of the full scan was assigned a microstate label. This backfitting was performed using previously described methods (Koenig et al., 1999; Michel et al., 2009). Each GFP peak in the full dataset was labelled as a state based on the microstate centroid map with minimum distance. All other samples were given the same state label as their nearest GFP peak.

For each condition (rest/task), the microstate maps were extracted only from the first MEG scan, but were backfit to both scans to provide validation of microstate statistics across recording sesisons.

### 2.2 Microstate statistics

We studied a number of spatiotemporal statistics of the resulting source MEG microstate sequences. Global statistics of the microstate sequences include GEV (Murray et al., 2008), mean duration of microstates (Koenig et al., 2002), Hurst exponent of the sequences (Van De Ville et al., 2010), and microstate complexity (Tait et al., 2020b). Microstate complexity values were normalized against its theoretical asymptotic upper bound (Lempel and Ziv, 1976; Liu et al., 2016; Zhang et al., 2016), to result in a measure *∈*(0, 1]. Class-specific statistics included mean duration of the microstates within a particular class (Koenig et al., 2002; Lehmann et al., 2005), coverage of a class (the percentage of time spent within a class) (Lehmann et al., 2005), and occurrences of the class (number of times the state appears per second) (Lehmann et al., 2005). We also calculated the Markov and syntax matrices (with and without self-transitions respectively) (Lehmann et al., 2005; Nishida et al., 2013; von Wegner et al., 2017), the information-theoretical zeroth and first order Markov statistics and their *p*-values (von Wegner et al., 2017), and tested for non-random microstate syntax (Lehmann et al., 2005; Nishida et al., 2013). Details and results of analysis to quantify non-stationary, i.e. by comparing microstate statistics against random fluctuations in a stationary process, are given in Supplementary Text S2.2.

### 2.3 Microstates to study dynamic functional connectivity

To derive functional connectivity for a given microstate, we adapt the EEG microstate segmented phase locking method of Hatz et al. (2015, 2016). The method is described in detail in Supplementary Text S1.4, and a visual overview of the pipeline is given in Supplementary Figure S1. Microstate analysis was first performed on broadband (1-30 Hz) data. We subsequently filtered the data into one of three narrow bands (theta 4-8 Hz, alpha 8-13 Hz, beta 13-30 Hz), and calculated the Hilbert transform. For a given microstate, all samples of the analytic signal within this microstate class were concatenated. Under the assumption that microstates are associated with specific patterns of phase synchronization (i.e. the hypothesis we wish to test), we subsequently epoched these concatenated samples into non-overlapping windows of 1280 samples (equivalent to 5-seconds), and calculated phase synchronization for each window using the weighted phase lag index (wPLI) (Vinck et al., 2011; Colclough et al., 2016). wPLI was chosen because it is invariant to instantaneous phase synchrony. This is advantageous in our study for two reasons. Firstly, our networks will not be influenced by leakage in the inverse algorithm. Secondly, since microstate analysis involves topographic clustering of the instantaneous patterns of activation, by studying only phase-lagged connectivity we are not simply quantifying the same effect as microstate analysis using a different method. This method was repeated for each microstate class, to obtain functional connectivity patterns for each class.

This methodology assumes that microstates are associated with a unique pattern of phase synchronization, and therefore it is meaningful to calculate a single functional network for all samples in a given class. To test this hypothesis, we applied multi-variate pattern analysis (MVPA) using the MVPA-Light toolbox (Treder, 2020).

Across all participants, resting-state scans, 1280-sample segments, and microstate classes there were 5413 networks per frequency, each associated with a single microstate class. The weighted degree distributions were used as features in a multi-class classifier for microstate label. Results were 5-fold cross-validated. Statistically significant classification accuracy (via permutation testing) is strongly suggestive that microstates are associated with unique patterns of phase synchronization, demonstrating that microstate class can be predicted given functional connectivity derived from any 1280 samples from that microstate class. Details on classification are given in Supplementary Text S1.4.

### 2.4 Analysis of task microstates

To compare microstates derived from resting-state and task conditions, we separately performed microstate analysis on data recorded from each condition and aligned microstates across conditions using a template matching algorithm. Microstate statistics were averaged over all data sets available within a condition for a participant. Microstate statistics between conditions were compared using paired Wilcoxon sign-rank tests.

To assess whether specific states were associated with auditory stimuli, we calculated an expected probability of all states under the null hypothesis of no association between state likelihood and stimuli by calculating the coverage of each state over samples in the pre-stimulus periods (up to 100 ms prior to the stimuli) across all task scans and all participants. For a given latency in the range 0-350 ms following the stimulus, we calculated an observed count of states by calculating the number of occurrences of each state over all stimuli and all participants. A *χ*^2^ test was used to compare the observed count with the expected count (equal to expected probability multiplied by total number of stimuli). To post-hoc test for deviations of the count of each microstate class from the expected count, we used the Pearson residual (Agresti, 2012), calculated as

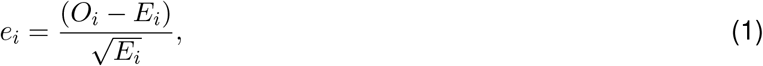

3where *e*_*i*_ are the residuals, and *O*_*i*_ and *E*_*i*_ and the observed and expected counts of microstate *i* respectively. The normalization by 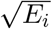 standardizes these scores to be distributed as the standard normal distribution under the null hypothesis. The *χ*^2^ statistics can be written as 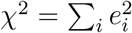, and hence microstates with large magnitude of *e*_*i*_ contribute to larger values of *χ*^2^.

### 2.5 Participants and Data

#### 2.5.1 Participants

Thirty healthy participants (16 female, 9 male) were recruited from Cardiff University School of Psychology participant panel (age range 17-28 years, median age 20 years). All participants had normal or corrected-to-normal vision, and none reported a history of neurological or psychiatric illness. Written consent was obtained from all participants. The study was approved by the Cardiff University School of Psychology Research Ethics Committee.

#### 2.5.2 MEG Acquisition and preprocessing

Whole-head MEG recordings were made using a 275-channel CTF radial gradiometer system (CTF Systems, Canada) at a sampling rate of 1200 Hz. An additional 29 reference channels were recorded for noise cancellation purposes and the primary sensors were analysed as synthetic third-order gradiometers (Vrba and Robinson, 2001). One sensor was turned off during recording due to excessive sensor noise (i.e., *N*_*x*_ = 274 gradiometers). Horizontal and vertical electro-oculograms (EOG) were recorded to monitor blinks and eye movements. The horizontal electrodes were placed on temples, and vertical ones, above and below the eye. For MEG/MRI co-registration, the head shape with the position of the coils was digitised using a Polhemus FASTRAK (Colchester, Vermont).

MEG was recorded over two sessions, recorded on separate days (1-24 days between sessions). In each session, eight minutes of continuous resting-state was recorded, for which participants were instructed to sit comfortably in the MEG chair while their head was supported with a chin rest and with eyes open focus on a fixation point on a grey background. Additionally, ten minutes of passive task activity was recorded in each session, for which participants were instructed to watch an emotionally neutral movie while periodic auditory stimuli (separated by 500 ms) were played through headphones. Half of the stimuli consisted of standard tones, while the other half were a mixture of duration, gap, direction, intensity, and frequency deviates.

For preprocessing, MEG data was imported to Fieldtrip (Oostenveld et al., 2011), bandpass filtered at 1-100 Hz (4th order two-pass Butterworth filter), notch filtered at 50 and 100 Hz to remove line noise, and downsampled to 256 Hz. Visual and cardiac artifacts were removed using ICA decomposition (aided by EOG recordings), using the ‘fastica’ algorithm (Hyvarinen, 1999). For both task and resting-state data, between 2 and 6 components were removed per scan (median 4 for rest, 3.5 for task).

#### 2.5.3 Source reconstruction

All participants also underwent a whole-brain MRI scan on a Siemens 3T Connectom MRI scanner and a 32-channel receiver head coil (Siemens Medical Systems). We used a T1-weighted magnetization prepared rapid gradient echo sequence (MPRAGE; echo time: 3.06 ms; repetition time: 2250 ms sequence, flip angle: 9*°*, field-of-view: = 256 *×* 256 mm, acquisition matrix: 256 *×*256, voxel size: 1*×* 1*×* 1 mm).

From the T1-weighted MRI image, extraction of the scalp, brain, and cortical surfaces was performed with Freesurfer (Dale et al., 1999). Vertices of the cortical surface were labelled according to the HCP230 atlas optimized for MEG studies (Tait et al., 2020a). The scalp surface was used to align the structural data with the MEG digitizers. A single shell volume conduction model (Nolte, 2003) was constructed in Fieldtrip using the brain surface (downsampled to 500 vertices). The cortical mesh was downsampled to approximately 10,000 vertices to generate a set of dipole locations using the ‘iso2mesh’ software (Fang and Boas, 2009), and dipoles were oriented normal to the cortical surface (Dale et al., 2000; Hillebrand and Barnes, 2003).

Source reconstruction used the eLORETA algorithm (Pascual-Marqui, 2007, 2009) implemented in Fieldtrip. Source data was parcellated using the HCP230 atlas labels (Tait et al., 2020a) by taking the time course of the first principal component of all voxels with an ROI. Source time courses were bandpass filtered in the 1-30 Hz frequency band, and head localization coils were used for offline head motion correction (Stolk et al., 2013).

### 2.6 Simulations

Simulations were performed to test the ability of the microstate methodology to estimate a ground truth microstate sequence and spatial maps, and compare modified against other methodologies for neural state identification such as PCA, ICA (Hyvarinen, 1999), and hidden Markov modelling (Vidaurre et al., 2016). Details of the simulations and settings for other algorithms are given in Supplementary Text S1.3. Artificial generative sequences were generated using a random walk decision tree approach, detailed in Supplementary Text S1.3, which reverse engineers a method to transform microstates to random walks (Van De Ville et al., 2010; von Wegner et al., 2016, 2018). Neural dynamics were then generated by assigning each microstate a Wilson-Cowan neural mass model of resting-state dynamics (Deco et al., 2009; Abeysuriya et al., 2018), and stimulating the oscillator associated to the active state. Finally, the states were assigned spatial maps of the same spatial dimensionality as our data, i.e. 230 ROIs. The choice of spatial map is arbitrary, so to increase neurophysiological realism we used open access resting-state networks derived from fMRI (Smith et al., 2009) aligned to our brain atlas. ROI time series were generated as a combination of a linear projection of each microstate’s neural dynamics onto the microstate maps and pink noise with SNR=1. Twenty repetitions of the simulations were performed, and simulations were repeated for *k* = 4 and *k* = 10 states. Four statistics were used to assess the quality of the microstate estimation against the ground truth, namely GEV, map similarity, temporal mutual information (MI) the ground truth and estimated sequence, and the normalized mutual information only at the GFP peaks (MIg).

It should be highlighted that these simulations do not aim to reproduce biophysical mechanisms of microstate generation as currently these mechanisms are not well understood. Due to this, some key properties of true microstate sequences such as spontaneous transitioning and associations between GFP and state stability (Koenig and Brandeis, 2016) are not recreated by our model. However, this should not crucially alter the outcome of our results, since the aim of the simulations was to generate artificial data with few underlying model assumptions and known ground truth maps/sequences.

### 2.7 Data and Code Availability

Data preprocessing and source reconstruction used Fieldtrip version 2019-07-16 (Oostenveld et al., 2011). Microstate analysis, simulations, and statistics used a set of custom scripts written in MATLAB R2017a, which will be made freely available but are currently available only upon request. Data used in this study are available upon request.

## 3 Results

### 3.1 Simulations

We used a neural-mass model to generate synthetic time courses of electrophysiological data with a known number of microstates. The simulations were then used to test the *k*-means microstate algorithm against the ground truth, and to compare the performance against using PCA, ICA, and HMM for estimating brain states (see Supplementary Text S1.3.1 for a description of the methods). Figure 1 shows the simulation pipeline and results of analysing the simulated datasets. Results of GEV, map similarity, and mutual information analyses were largely consistent between simulations of 4 microstates and 10 microstates. In terms of GEV (Figure 1D), the three highest performing methodologies were *k*-means applied to amplitude envelopes, HMM applied to amplitude envelopes, and *k*-means applied to source time courses. The remaining algorithms -PCA, ICA, and source HMM - performed notably worse.

**Figure 1.**
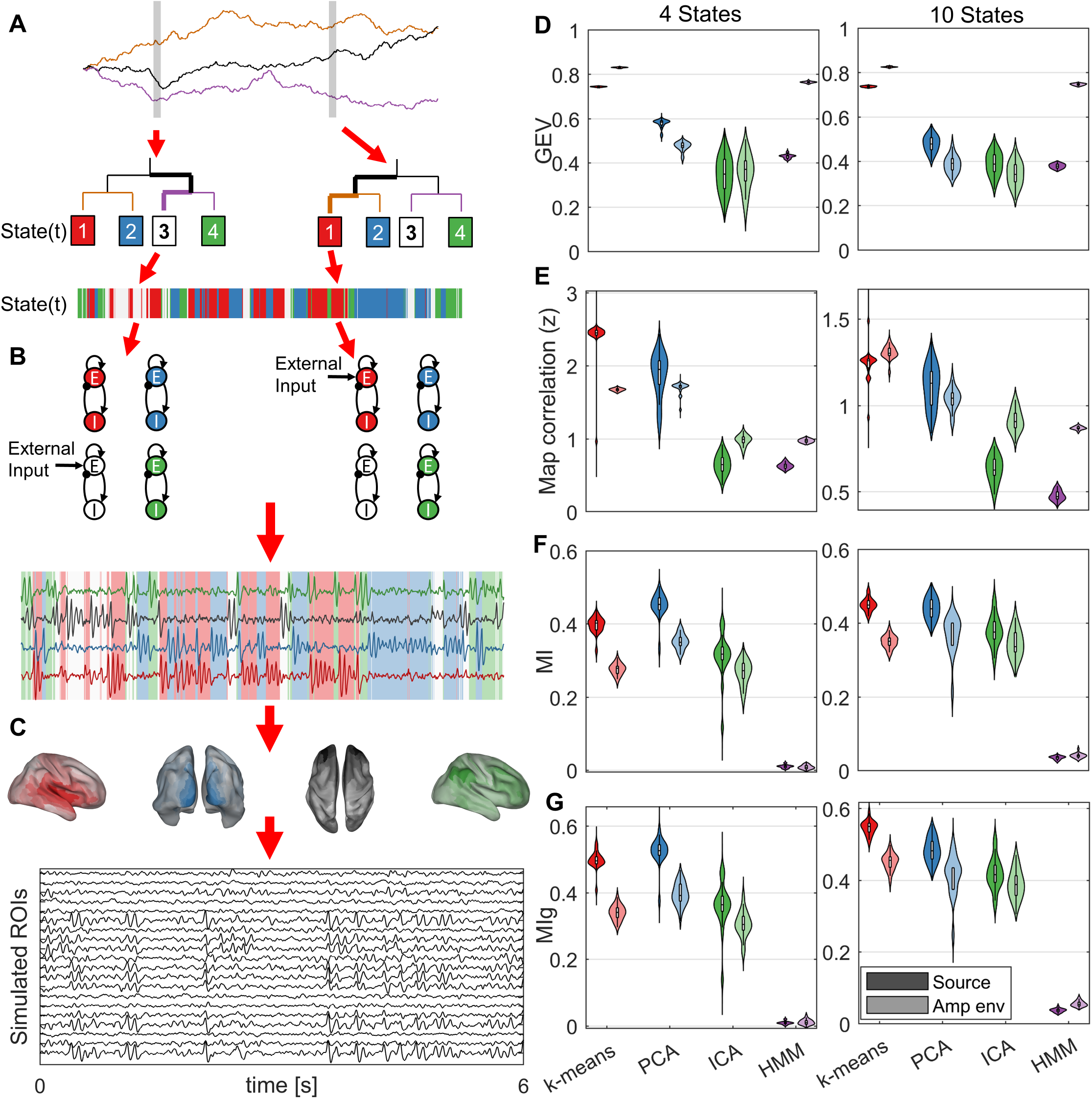
Simulating microstates. (A-C) Methodology for simulating microstates. Detailed description is given in Supplementary Text S1.3. (A) Generative sequences are simulated using a decision tree, the state at a given time (bottom; colours correspond to state) is decided by a set of random-walks (top) and a decision tree (middle). Colours of branches on the decision tree correspond to the colours of the random walks. The left branch is chosen if the walk is increasing, the right branch is chosen if the walk is decreasing. (Bottom) Each sample is assigned a state using this procedure. (B) Neural dynamics of microstates are simulated by assigning each state an neural mass model (NMM). The NMMs receive constant input (*P*) at all times, and the NMM corresponding to the current active state receives additional input (*P*_*ms*_). The bottom shows simulated state time series, with background colour corresponding to active state. (C) Each state is assigned a spatial map and the time courses are linearly mixed according to this map to generate simulated MEG/EEG/source data. Pink noise is added to this data. (D-G) Assessment of microstate sequences estimated from simulated data vs ground truth. (D) GEV, (E) correlation between estimated maps and ground truth maps, (F) mutual information between estimated sequences and ground truth sequences.

*k*-means and PCA most accurately recreated the ground truth maps in both source and amplitude domains, with *k*-means consistently outperforming PCA in all conditions except 4-state amplitude envelopes (Figure 1E). In terms of mutual information between ground truth and estimated sequences, the same two algorithms were also the highest performing (Figure 1F-G). In all conditions, source data outperformed amplitude data. Interestingly, the HMM had normalized MI of less than 5% for all simulations over all conditions (full data or GFP peaks, and amplitude or source), so performed poorly at recreating the ground truth sequences. As a consequence of these results, we used the source data as opposed to amplitude envelope data for all subsequent microstate analysis.

### 3.2 Resting-state microstate maps

We source-reconstructed MEG resting-state data from 30 participants.Across all participants, microstates in resting-state data were calculated from a sample of 150,000 GFP peaks using the *k*-means clustering algorithm. Figure 2A shows the GEV across these 150,000 peaks as *k* is varied from 2-40 states. The kneedle algorithm indicated that 10 states were optimum, and hence we proceeded to back-fit the results of the 10-state clustering to the full MEG scans. Figure 2B shows the GEV in the full datasets. In the first set of scans, from which the GFP peaks were sampled to estimate microstates, 10 states had a GEV of 63.97 *±* 0.64%. To ensure the states were generalizable and reproducible, we also back-fit the maps to a second independent scan from each participant, which was performed on a separate day and not used in the clustering analysis. There was a significant but small decrease in GEV in the second scan (62.13 *±*0.39%, *p* = 0.0082, Wilcoxon sign-rank test), as expected in cross-validation. Furthermore, multiple runs of the algorithm demonstrated high reproducibility (Supplementary Text S2.1).

**Figure 2.**
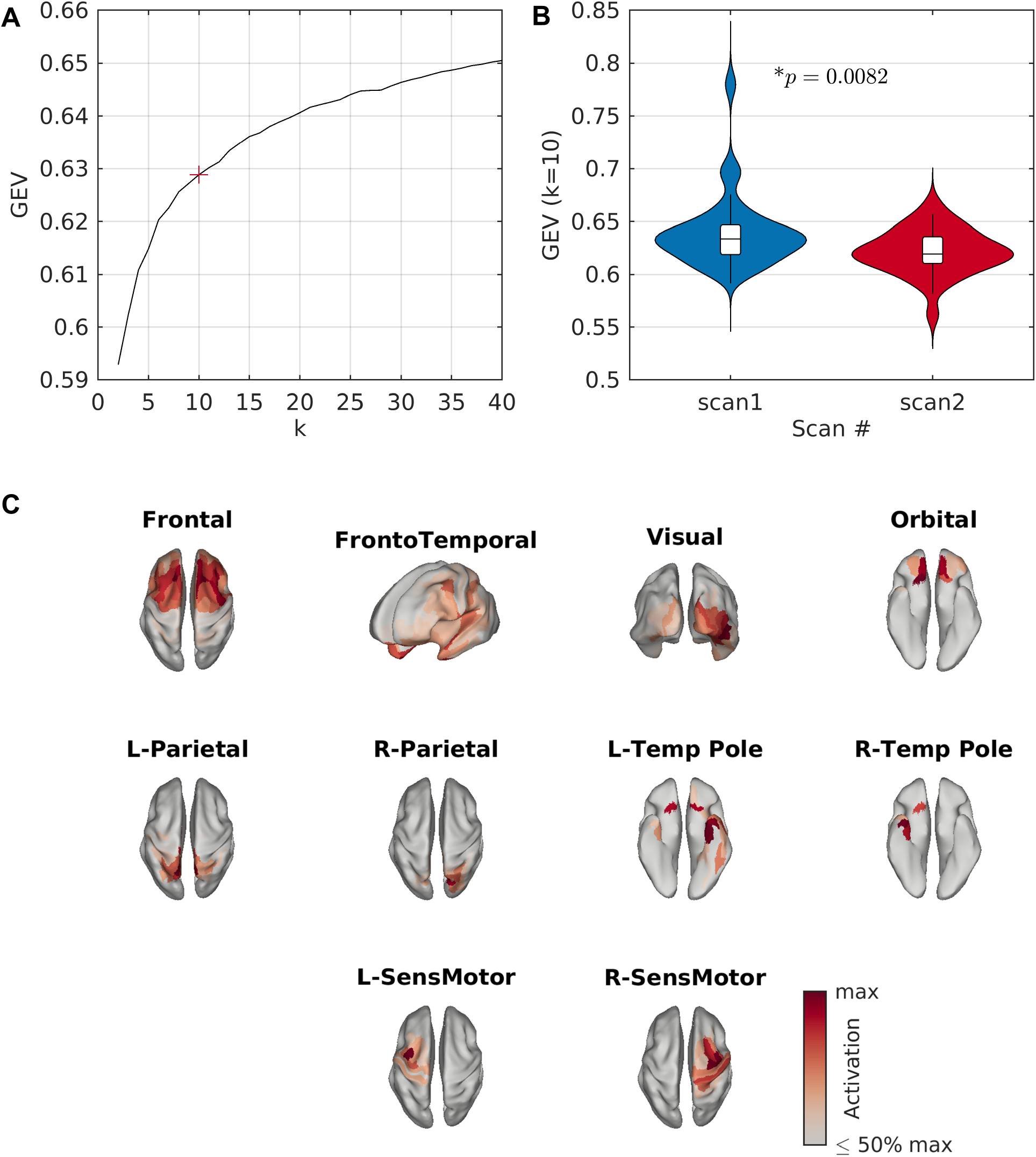
Resting-state microstates. (A) GEV vs number of states (*k*) for resting-state data. Here, GEV is calculated across the 150,000 peaks used for clustering. The kneedle algorithm found that *k* = 10 states was optimum, marked by a red ‘+’. (B) GEV across the full MEG scans, for *k* = 10. Distributions are shown across participants. Only GFP peaks from scan 1 for each participant were used for clustering, and hence scan 2 can be viewed as a replication/validation cohort. There is a small but significant decrease in GEV for scan 2. (C) Resting-state microstate maps derived from the *k*-means clustering algorithm for *k* = 10 states.

Figure 2C shows the spatial map of each empirical MEG source-level microstate. Four bilateral maps were identified, including the frontal cortices, the fronto-temporal network, the visual cortex, and the orbital cortex. The remaining six maps could be grouped into three pairs of lateralized networks including medial/superior parietal, temporal poles, and sensorimotor networks.

### 3.3 Statistics of resting-state microstate sequences

We subsequently analysed the statistics of the estimated microstate sequences, reported in full in Supplementary Table S3. Microstates had a mean duration (MD) across all classes of 59.86 *±* 1.09 ms (Supplementary Table S3). This was significantly longer than could be explained by random fluctuations in a stationary Gaussian process with the same power spectra, cross-spectra (and covariances), and distribution of data (Supplementary Text S2.2), suggestive of non-stationarity and the existence of stable microstates in empirical resting-state MEG. Durations of each state individually are shown in Figure 3.

**Figure 3.**
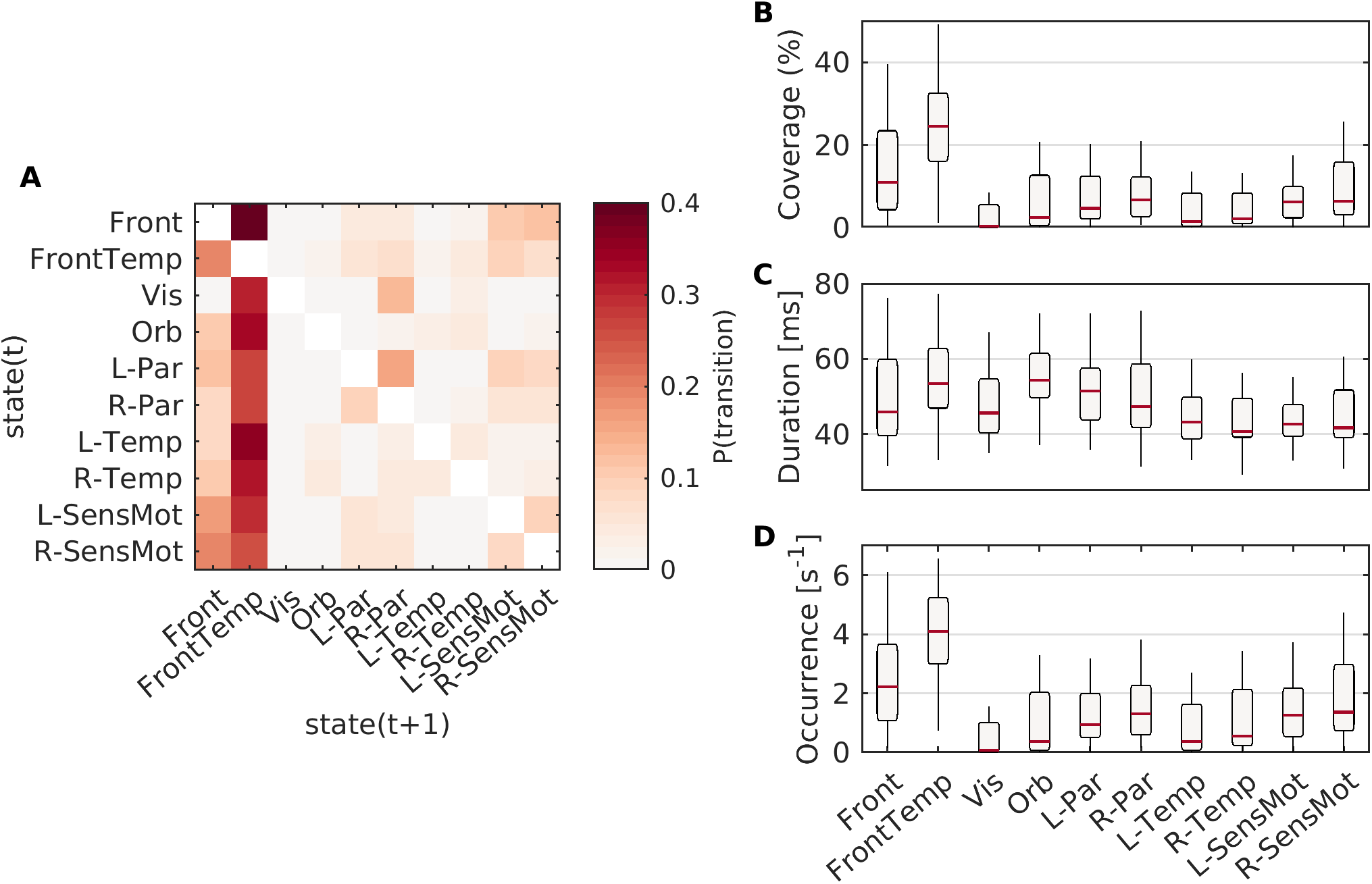
Syntax of resting-state microstates. (A) The average syntax matrix from resting-state data. (B-D) Coverage, duration, and occurrence of each of the ten resting-state microstate maps.

Of the 10 microstates, the frontal and frontotemporal states had the highest coverage and highest number of occurrences per second (Figure 3). This can be observed in the group-level syntax matrix (Figure 3A), in which almost all states tend to transition most predominantly to the frontotemporal state, which in turn is most likely to transition to the frontal state. However, random sampling of states with heterogeneous occurrences per second is not sufficient to explain the syntax matrix, as the syntax permutation test (Lehmann et al., 2005) demonstrated significant differences between the group level matrix and the matrix expected due to state probability (*p* =4 *×* 10^*−*4^). Similarly, when accounting for duration and coverage (i.e. including self transitions in the Markov matrix), information theoretical analysis still demonstrated that the zero’th order Markov (*G*_0_) property was, for all participants, far greater than expected under the null hypothesis of random sampling based on coverage of states (Supplementary Table S3). The transitioning matrix could also not be explained by a first order Markov process, as demonstrated by the Markov *G*_1_ information theoretical analysis (Supplementary Table S3). Combined, these results suggest structured and non-Markovian transitioning in the microstate sequence.

The structured, non-stationary and non-Markovian nature of microstate transitions was additionally supported by global statistics. The Hurst exponent (H) was approximately 0.68*±* 0.01, suggestive of long range temporal correlations. The normalized microstate complexity (C) was 0.60 *±* 0.01, suggesting a complex sequence which strikes a balance between highly repetitive and ordered (C approaching zero) and random sampling (C approaching one). These global statistics were significantly different than expected from random fluctuations in a stationary process (Supplementary Text S2.2), further evidencing the non-stationary nature of resting-state MEG.

### 3.4 Microstate-specific functional connectivity

Our next aim was to test the hypothesis that different microstate classes are associated with distinct patterns of functional connectivity in the brain, and reflect the rapid transitioning in dynamic phase synchronization patterns. To do so, we used multi-class MVPA to test whether microstates could be predicted from microstate-segmented wPLI connectivity matrices (see section 2.3). Table 3 shows the classification accuracy and permutation testing *p*-values for each frequency band, while Figure 4A shows the confusion matrices for this classification.

**Table 3:** MVPA classification statistics for microstate-segmented connectivity. Accuracy is given as a mean and standard error across five repetitions.

| Band | Accuracy | *p*-value |
| --- | --- | --- |
| Theta | $0.2068 \pm 0.0084$ | $< 10^{-3}$ |
| Alpha | $0.3281 \pm 0.0120$ | $< 10^{-3}$ |
| Beta | $0.3083 \pm 0.0112$ | $< 10^{-3}$ |

**Figure 4.**
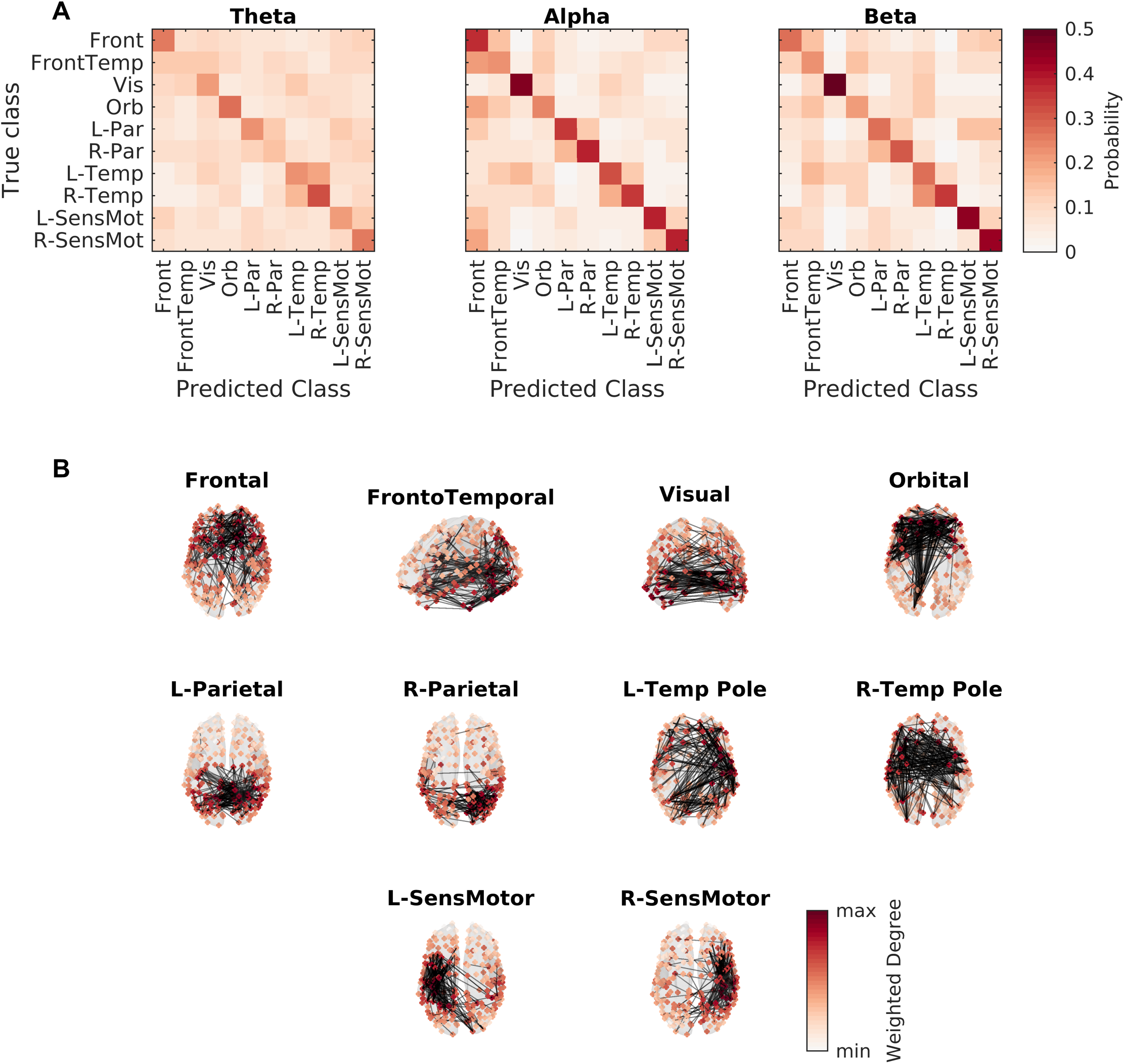
Microstates are associated with specific patterns of phase synchronization. (A) Confusion matrices for the MVPA analysis, for three frequency bands used for functional connectivity calculation. Cell *i, j* corresponds to the probability the classifier would predict class *j* given a microstate label *i*. (B) Functional connectivity patterns. Node colours show degree distributions (used for classification in the MVPA analysis), while edges show the top 1% of edges that deviate from static background connectivity (see Supplementary Text S1.4 for details).

The classification accuracy was significantly above the chance level of 10% (*p <* 10^*−*3^, 1,000 permutation tests from 1,000 surrogates), suggesting distinct functional connectivity patterns among microstates. The confusion matrices demonstrate interesting patterns. In the alpha and beta bands, the visual microstate is the one most accurately predicted by the connectivity matrix, while in theta the temporal microstates were more accurate. Unsurprisingly because of their overlap, frontal, frontotemporal, and orbital networks had reasonable degrees of confusion in all bands. In the alpha band, the frontal network was also sometimes confused with bilateral sensorimotor networks.

Furthermore, lateralized microstates were regularly confused with their counterparts in the opposite hemisphere, suggesting that while activation patterns in these states may be lateralized, the states are parts of a larger interhemispherical network. Indeed, Figure 4B shows the alpha band network structures, and it is clear that the functional connectivity structure for both lateralized parietal and temporal microstates demonstrated many interhemispherical connections.

### 3.5 Microstates of auditory evoked response

Finally, we aimed to test whether source-level MEG microstates vary between rest and task. To do so, we re-ran the microstate clustering analysis on MEG data during a passive auditory task. The optimum number of microstates in the task data was *k* = 9. This was one fewer than at rest, yet there were no significant differences in the GEV between nine states in task and ten states at rest (Figure 5A-B; *p* = 0.9099).

**Figure 5.**
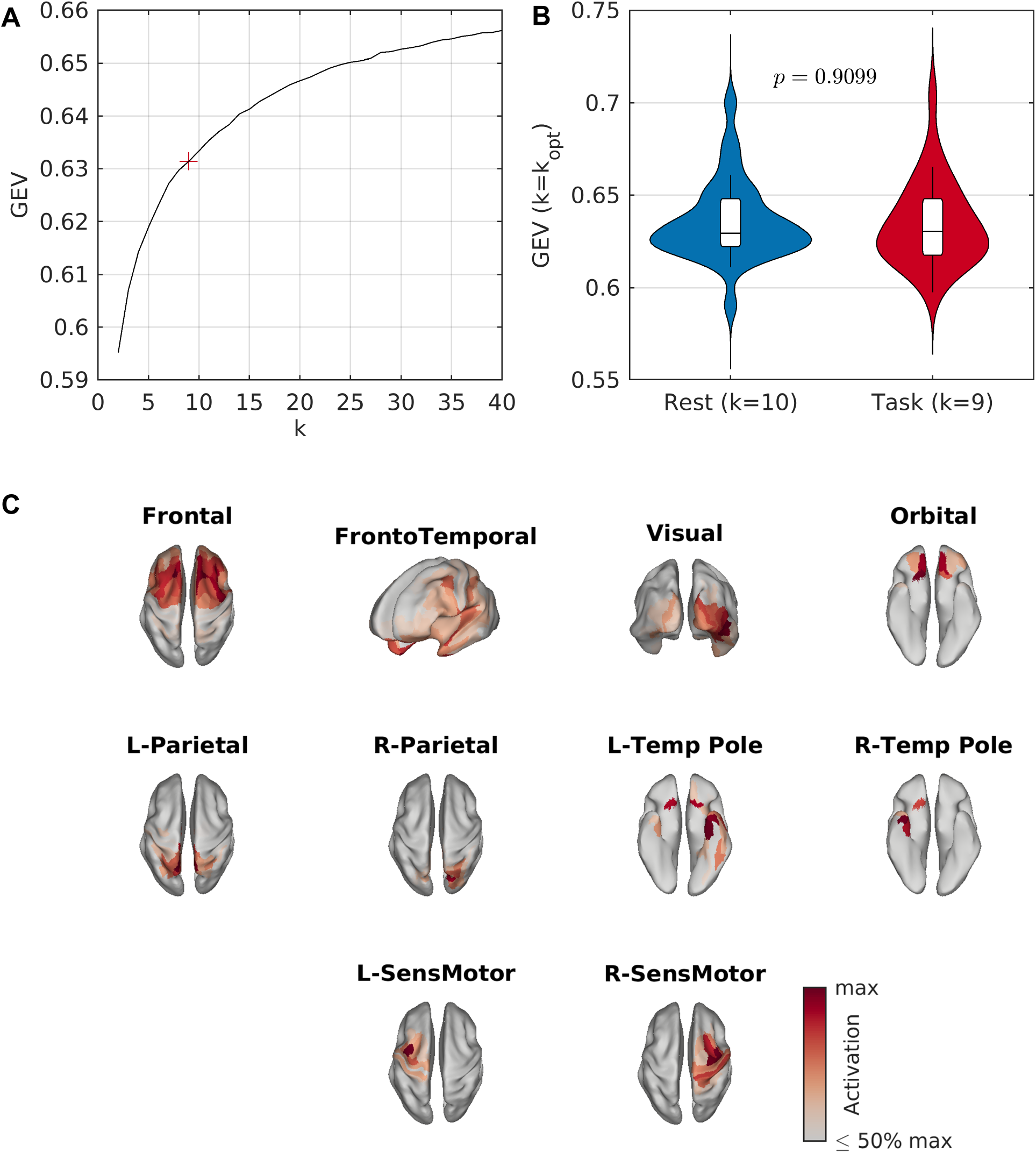
Auditory task microstates. (A) GEV vs number of states (*k*) for resting-state data. Here, GEV is calculated across the 150,000 peaks used for clustering. The kneedle algorithm found that *k* = 10 states was optimum, marked by a red ‘+’. (B) GEV across the full MEG scans, for *k* = 10. Distributions are shown across participants. Only GFP peaks from scan 1 for each participant were used for clustering, and hence scan 2 can be viewed as a replication/validation cohort. There is a small but significant decrease in GEV for scan 2. (C) Resting-state microstate maps derived from the *k*-means clustering algorithm for *k* = 10 states.

Maps for the nine states derived from the task data are shown in Figure 5C. The task-based states closely corresponded to those from rest. The two lateralized medial parietal states in the resting-state data were represented by a single medial parietal state in the task data, explaining the presence of one fewer state. The frontal and visual states demonstrated differences between conditions, with the frontal state appearing more right lateralized and the visual state having greater spatial extent. All other states closely recreated their resting-state counterparts. There were no significant differences in global statistics of the microstate sequences between rest and task, including mean duration (*p* = 0.5170), Hurst exponent (*p* = 0.2452), or complexity (*p* = 0.8774).

To uncover whether any states were time-locked to auditory stimuli, we used *χ*^2^ tests to compare pre-stimulus and post-stimulus coverage of each state. The time evolution of *χ*^2^ is shown in Figure 6A. There is a significant difference in the observed and expected microstate counts between 70 and 280 ms following the stimulus, peaking at 113 ms. This is in line with the auditory N100 evoked response. The differences in microstate count are largely driven by a significantly increased coverage of the frontotemporal state, with resulting decreased coverage of several other microstates including a significantly decreased coverage of the medial parietal state (Figure 6B). Since the frontotemporal state includes the auditory network (Figure 5C), these results are in line with increased activation of the auditory network as a response to an auditory stimulus.

**Figure 6.**
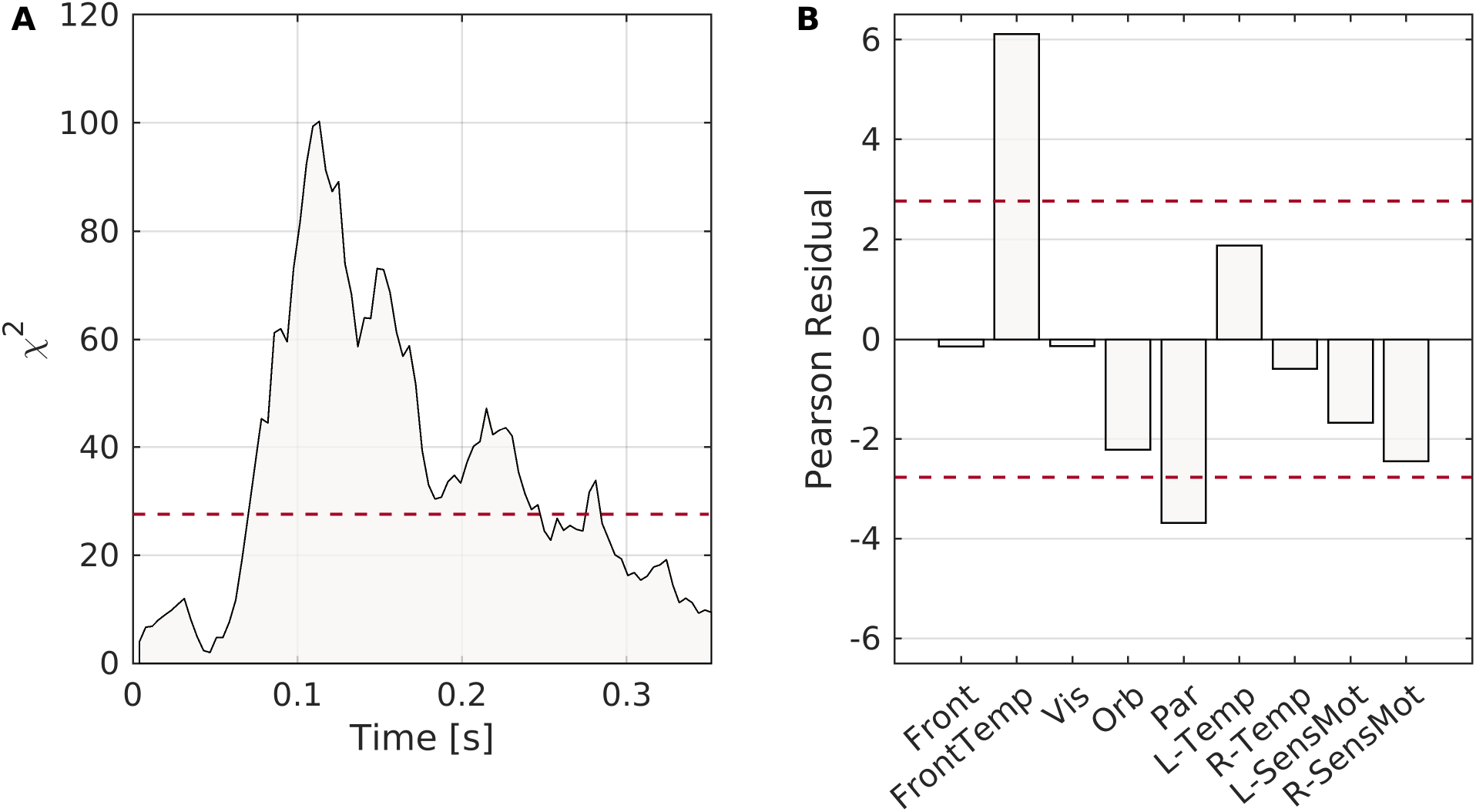
Microstate response to auditory stimuli. (A) *χ*^2^ distance between observed and expected counts of each microstate vs time after stimulus. The peak response was 113 ms after the stimulus. (B) Pearson residuals for each state. For large sample sizes, under the null hypothesis the Pearson residual is approximately normally distributed. Here, Pearson residuals are averaged across all samples around the peak with a value of *χ*^2^ greater than one-half the peak value. For both plots, dotted red lines show the Bonferroni corrected 95% significance level. Hence, these results suggest that between 70-280 ms following the stimulus the microstate response is significantly different that expected, due to a significantly increased likelihood of the frontotemporal state and decreased likelihood of the parietal state.

## 4 Discussion

Understanding the mechanisms underpinning discrete brain state generation is currently a leading question in the field of neuroscience, which may aid with understanding behaviour and cognition (Cohen, 2018) and neurological diseases (Khanna et al., 2015). One approach to do this in sensor EEG is microstate analysis (Michel and Koenig, 2018), but this approach cannot be directly applied to source data. In this study, we have presented a modified source-space microstate pipeline and demonstrated the validity and robustness of source-space MEG microstates. Using a *k*-means clustering algorithm, we showed that 10 microstates can be reliably estimated from resting-state MEG data, and their spatial distributions are similar to those quantified from task-based data. Source-space MEG microstates vary in their statistics such as coverage and occurrence probabilities, and are associated with distinct functional connectivity signatures. These results support the use of MEG microstates to understand dynamic functional connectivity during rest and task.

Interestingly, in resting-state data we observed the optimum number of states to be 10, explaining approximately 63% of the variance of the data. In resting-state sensor EEG, only four microstates have repeatedly been reproduced and observed to be the optimum, explaining approximate 60-85% of variance in the data (Michel and Koenig, 2018). There are a number of potential explanations for the differences between these results. One possible explanation is that EEG usually has lower spatial resolution than MEG, due to (typically) fewer sensors and a blurring of the electric field by the skull tissue. In a study with high-density EEG (204 electrodes), seven maps were optimum (Custo et al., 2017), supporting the hypothesis of higher spatial resolution potentially increasing the optimum number of maps. Another possible explanation is the criterion used for selecting the optimum number of maps, as many criteria are available in the literature (Michel and Koenig, 2018). Discussion of this point, including motivation for our choice of criterion, are given in Supplementary Text S1.2. The larger number of states in this study may also be due to performing clustering in source space as opposed to sensor space. There is a non-uniform distribution of signal-to-noise ratio (SNR) in the forward projection of the source dynamics to sensor space, and hence it is possible that states originating from low-SNR areas of the cortex or may be under-represented in the sensor space maps. Additionally, due to finite spatial sampling of M/EEG sensors, multiple source topographies may result in non-unique maps, and therefore multiple source-space states may appear similar at the sensor level. Both of these effects potentially result in under-estimation of the number of states at the sensor level. In HMM studies of source-reconstructed resting-state MEG, Baker et al. (2014) studied eight states while Vidaurre et al. (2018) found twelve to be optimum, while fMRI results have suggested ten reproducible resting-state networks (Smith et al., 2009) or 7-8 states based on *k*-means clustering of dynamic resting-state functional connectivity patterns (Allen et al., 2014), in line with our results demonstrating greater than four states at the cortical level.

The ten resting-state microstates included four bilateral networks and three pairs of symmetric lateralized networks (Figure 2). The cognitive relevance of these microstates should be a focus of future work. One approach to uncover the associations between microstates and cognition is to quantitatively compare microstate statistics between cognitive states. Such approaches have widely been performed in the sensor-space EEG microstate literature to gain insight into the functional significance of brain microstates (Brodbeck et al., 2012; Britz et al., 2014; Milz et al., 2016; Seitzman et al., 2017; Zappasodi et al., 2019), and the work presented here opens new pathways to gain deeper anatomical insight at the cortical level. Here, we demonstrated that statistics of our source-space cortical microstates differed between rest and a passive auditory mismatch paradigm (Figure 5-6). We found hyperactivity of the frontotemporal microstate, which includes the auditory cortex, approximately 100ms following an auditory stimulus. Interestingly, the well-studied auditory evoked response has been localized to the the auditory cortex and first peaks around 100ms following the stimulus (Picton, 2010), known as the N100 response, in line with our microstate results. Future work should build upon these approaches to examine the functional significance of these states.

A complementary approach is to uncover associations between microstates and the well studied resting-state networks (RSNs) widely studied in fMRI, which have been associated with cognitive domains in large cohort studies (Smith et al., 2009). In sensor-space EEG, studies have demonstrated associations between microstates and fMRI-RSNS through convolution of the microstate time courses with a haemodynamic response function and general linear modelling (Britz et al., 2010; Musso et al., 2010; Van De Ville et al., 2010; Yuan et al., 2012; Abreu et al., 2020; Xu et al., 2020; Zoubi et al., 2020), or through correlations between RSNs and microstate statistics (Schumacher et al., 2019). An advantage of working in source space for this purpose is that spatial patterns of microstate activations can be associated with RSNs when directly comparable brain atlases are used for parcellation of the brain dynamics in the M/EEG and fMRI data.

While the activation patterns of our microstates did not directly correspond to the well known resting-state networks (RSNs) often reported in fMRI studies (Smith et al., 2009), interesting insight into this relationship could be gained by studying spatial patterns of synchrony associated with each microstate. Through the use of machine learning - specifically MVPA (Treder, 2020) - we found a significant association between active microstate class and cortical patterns of phase synchronization (Figure 4, Table 3). The spatial patterns of synchrony did not directly reflect the associated microstate maps, and the functional connectivity patterns identified here may give insight into the relationship between microstates and RSNs. A key example is the default mode network, which contains the medial/orbital frontal, medial parietal, and lateral parietal regions (Smith et al., 2009). No single microstate had an activation pattern containing all of these ROIs, yet phase-locking patterns indicted microstates demonstraing medial/orbital frontal to medial parietal connectivity and medial parietal to lateral parietal connectivity (Figure 4B). Future work should involve simultaneous high-density source-reconstructed EEG and fMRI to study the relationship between source-space microstates and RSNs.

The microstate connectivity approach presented here also has potential as an alternative approach for studying dynamic functional connectivity. A typical approach for dynamic functional connectivity in the current literature is the use of a sliding window (de Pasquale et al., 2010; Brookes et al., 2014; O’Neill et al., 2015; de Pasquale et al., 2016; Lopes et al., 2020), which is limited by the arbitrary choice of window size meaning development of novel methods beyond the sliding window are crucial (O’Neill et al., 2018). Our MVPA analysis indicated that windowing via microstate labelling and concatenation of samples within a state (i.e. microstate-segmented functional connectivity (Hatz et al., 2015, 2016)) is a powerful option for the study of dynamic functional connectivity states at a fast time scale without setting an *a priori* window length. However, this approach has some limitations. While MVPA classification was significant for all frequency bands, accuracy was not perfect and there may be overlaps between functional connectivity patterns of lateralized states (Figure 4). A second limitation of this approach is that it relies on the assumption of a discrete number of functional connectivity states. While this assumption does not apply in general to sliding-window connectivity studies, common subsequent analyses such as clustering networks (Allen et al., 2014; O’Neill et al., 2015; Mheich et al., 2015; Hassan et al., 2015) or recurrence analysis (Lopes et al., 2020) make similar assumptions, and hence microstate-windowing can be viewed as an alternative to these approaches without the reliance on window length.

In recent years several alternative methodologies have been proposed for source-space brain state estimation based on Hidden Markov Models (HMMs) (Baker et al., 2014; Gä rtner et al., 2015; Vidaurre et al., 2016, 2018; Taghia et al., 2018). The use of a microstate pipeline instead of HMMs in this study was motivated by evidence suggesting that for electrophysiological resting-state data, the underpinning assumptions of HMMs for brain states are not always met. Koenig and Brandeis (2016) highlighted that HMMs cannot capture the empirical relationship between GFP and state stability, while Gschwind et al. (2015) discussed the importance of long-range dependencies in the microstate sequence which are not accounted for by HMMs. Conversely, the source-space microstate pipeline presented here is based on data-driven clustering and hence does not rely on many specific assumptions to be met - a detailed discussion of the assumptions and their justification for electrophysiological data is given by Michel and Koenig (2018) - and hence may be more applicable to a wider range of datasets. This was demonstrated by our simulations. Simulated data was generated through stimulation of neural mass models, with stimulation patterns following a non-Markovian sequence. Hence, our biophysically informed model did not meet the assumptions of the HMM. In these simulations, we found the microstate pipeline greatly outperformed the HMM approach in terms of estimate the ground truth maps and brain-state sequences (Figure 1). It should however be highlighted that the mechanisms underpinning the physiological origins of microstates in the human brain are currently not well understood and may not be reflected by our simulations, so these results should not be interpreted as evidence that HMMs perform poorly for real resting-state brain data. Future work should include simulations under a range of models and assumptions and a deeper analysis of various task-related data sets in order to perform a more detailed comparison between clustering approaches and identify the correct circumstances under which each approach should be used.

### 4.1 Conclusions

We have presented a source-space microstate pipeline for estimating electrophysiological brain states. We uncovered ten resting microstates which were associated with distinct patterns of activation and phase synchrony across the cortex, and demonstrated these microstates were the result of stable non-stationary states arising as opposed to random fluctuations in a stationary process. The microstates were associated with cognitive state; in particular the resulting microstate probabilities were altered as a response to auditory stimulus, driven by hyperactivity of the frontotemporal microstate. Our results suggest the methodology presented here is a powerful tool for studying anatomically interpretable brain states and dynamic functional connectivity at the millisecond scale.

