## Supplementary for "MEG cortical microstates: spatiotemporal characteristics, dynamic functional connectivity and stimulus-evoked responses"

#### S1 Supplementary Methods

##### S1.1 Mathematical description of the generalized microstate-pipeline

Let $X \in \mathbb{R}^{N\times T}$ be an electrophysiological data set, where $N$ is the spatial dimensionality (number of M/EEG sensors, or source space dipoles/ROIs) and $T$ is the number of time samples. Each column of $X$ is a spatial map, i.e. a sample of data $\mathbf{x}(t) \in \mathbb{R}^{N\times 1}$. Then the aim of microstate analysis is to identify a set of spatial maps with high signal-to-noise ratio (SNR) and cluster these maps into a small number of discrete states (Pascual-Marqui et al., 1995; Koenig et al., 1999; Khanna et al., 2015; Michel and Koenig, 2018). The most widely used methodology for EEG is the modified k-means algorithm presented by Pascual-Marqui et al. (1995) and Koenig et al. (1999), although a range of methods exist (Michel and Koenig, 2018) which have been demonstrated to give consistent spatial topographies (Khanna et al., 2014) and information-theoretically invariant temporal sequences (von Wegner et al., 2018). There are steps in the EEG microstate methodology which limit application to MEG sensor space or M/EEG source space data (outlined in Main Text Section 2.1), and to overcome this we here present the EEG microstate $k$-means algorithm (Pascual-Marqui et al., 1995; Koenig et al., 1999) in a framework which allows for generalization to other large scale electrophysiological datasets such as M/EEG sensor and source space based on an appropriate modality-specific transformation of the each spatial map $\mathbf{y} = f(\mathbf{x})$.

In this generalized framework, the EEG algorithm (Pascual-Marqui et al., 1995; Koenig et al., 1999) is recovered by re-referencing the data to a common average, i.e. using the transformation $\mathbf{y} = \mathbf{x} - \mu_{\mathbf{x}}$, where $\mu_{\mathbf{x}}$ is the average of $\mathbf{x}$ over sensors. The equivalent transformations for other modalities are given in Table S1. For sensor MEG, no transformation of the data is required (including no re-referencing). For source space, where source flipping may be an issue, there are two options. One is to use the amplitude envelope of the data (row labelled “Amplitude” in Table S1). However, this requires a narrow band signal and rejects phase information. A second option is to simply take the (element-wise) absolute value of the samples (row labelled “Source” in Table S1), which can be applied to broadband data and mostly maintains phase information (excluding phase differences of $\pi$).

| Modality | Spatial transformation |
|---|---|
| EEG | $\mathbf{y} = \mathbf{x} - \mu_{\mathbf{x}}$ |
| MEG | $\mathbf{y} = \mathbf{x}$ |
| Source | $\mathbf{y} = (\lvert x_1\rvert, \lvert x_2\rvert, \ldots, \lvert x_N\rvert)^T$ |
| Amplitude | $\mathbf{y} = (\lvert H(x_1)\rvert, \lvert H(x_2)\rvert, \ldots, \lvert H(x_3)\rvert)^T$ |

Supplementary Table S1: **Transformations used in generalized formulation of clustering analysis for each modality.** Here, $\mathbf{x} \in \mathbb{R}^{N\times 1}$ is a spatial topography with spatial dimension $N$ (i.e. a single time sample of multivariate data) with elements $(\mathbf{x})_n = x_n$, $\mu_{\mathbf{x}}$ is the mean of $\mathbf{x}$, $\lvert \cdot \rvert$ denotes absolute value, and $H(\cdot)$ denotes the (temporal) Hilbert transform (therefore $\lvert H(\cdot)\rvert$ is the instantaneous amplitude envelope).

The first step of the pipeline involves extracting samples with local peaks of SNR and map stability. These samples are peaks of the *global field power* (GFP), defined at a given time within our generalized framework as

$$\sigma(t) = \sqrt{\frac{1}{N-1}}\|\mathbf{y}(t)\| = \sqrt{\frac{1}{N-1}\sum_{n=1}^{N} y_n(t)^2} \tag{1}$$

The GFP is therefore the total deviation of the transformed map $\mathbf{y}$ from zero across space. For EEG, in which $y_n = x_n - \mu_{\mathbf{x}}$, this is equal to the standard deviation of the map, i.e. the total variation from the common average reference. In MEG and source space, the function is more appropriately viewed as a normalized vector norm of the map, and is therefore the total variation from zero field/current. Peaks of the GFP were here found using Matlab's findpeaks function. Appropriate pre-processing of the data including bandpass filtering is important here to generate a smooth GFP signal for accurate identification of peaks.

Once the set of GFP peaks has been extracted, we cluster the set of transformed spatial maps at these peaks into $k$ clusters. The $k$-means algorithm is outlined in Table 2 of the main text, and a clear graphical representation is given in Koenig et al. (1999) and Michel et al. (2009). The number of states $k$ is a free parameter, and must be chosen appropriately for the data. The algorithm initializes (step #1 in Main Text Table 2) by randomly selecting $k$ transformed maps as initial state centroids $\mathbf{C} \in \mathbb{R}^{N \times k}$. Here, we apply the $k$-means++ method of selecting initial maps (Arthur and Vassilvitskii, 2007; Tait et al., 2020), which has better performance and faster convergence than uniform sampling (Arthur and Vassilvitskii, 2007) which has commonly been used for microstate studies (Pascual-Marqui et al., 1995; Koenig et al., 1999; Michel and Koenig, 2018).

The subsequent steps of the algorithm are iterated until some convergence critera are met. Firstly, the distance between each transformed map and each cluster centroid is calculated (step #2 in Main Text Table 2). It is convenient to first define the *similarity* between a (transformed) map $\mathbf{y}$ and a centroid $\mathbf{c}$ (i.e. a column of $\mathbf{C}$) as

$$R(\mathbf{y}, \mathbf{c}) = \frac{|\mathbf{y}^T \cdot \mathbf{c}|}{\sqrt{\mathbf{y}^T \cdot \mathbf{y}}\sqrt{\mathbf{c}^T \cdot \mathbf{c}}}. \tag{2}$$

This equation is often called the *cosine similarity*, as it can equivalently be written as

$$R(\mathbf{y}, \mathbf{c}) = |\cos(\theta(\mathbf{y}, \mathbf{c}))|, \tag{3}$$

where $\theta(\mathbf{y}, \mathbf{c})$ is the angle between the two vectors $\mathbf{y}$ and $\mathbf{c}$, and hence it is clear that this value is in the range zero (orthogonal maps) to one (maps point in the exact same direction, ignoring polarity).

Subsequently, we can define the *distance* between maps as

$$D(\mathbf{y}, \mathbf{c}) = 1 - R(\mathbf{y}, \mathbf{c}) \tag{4}$$

For EEG, where both $\mathbf{y}$ and $\mathbf{c}$ have zero mean, it is clear from Equation 2 that the similarity is the absolute value of correlation, in line with the classic microstate $k$-means clustering algorithm (Pascual-Marqui et al., 1995; Koenig et al., 1999). For MEG and source data this can more readily be interpreted as the cosine similarity, i.e. the absolute value of the cosine of the angles between the vectors $\mathbf{y}$ and $\mathbf{c}$. For all modalities, $0 \leq D \leq 1$, where a value of one represents uncorrelated/orthogonal spatial maps, while a value of zero represents identical maps up to a scaling factor, i.e. $D(\mathbf{y}, \mathbf{c}) = 0 \iff \mathbf{y} = \alpha\mathbf{c}$, where $\alpha \in \{\mathbb{R} \setminus 0\}$.

Each map is subsequently labelled as belonging to the cluster with which it has minimal distance (or maximal similarity) to the cluster centroid (step #3 in Main Text Table 2), i.e.

$$L_i = \min_j(D(\mathbf{y}_i, \mathbf{c}_j)) \equiv \max_j(R(\mathbf{y}_i, \mathbf{c}_j)), \tag{5}$$

where subscript $i$ represents the $i$'th GFP peak with map $\mathbf{y}_i$, $j$ represents the $j$'th cluster with centroid $\mathbf{c}_j$, and $L_i \in \{1, 2, \ldots, k\}$ is the label assigned to peak $i$.

The centroid for each cluster is then updated (step #4 in Main Text Table 2). For a given cluster $j$ containing $M$ maps, let us define the matrix $\mathbf{Y}_j \in \mathbb{R}^{N \times M}$ such that each column of $\mathbf{S}_j$ is one of the transformed maps $\mathbf{y}$ in the cluster. As in the algorithm of Pascual-Marqui et al. (1995), the updated centroid of cluster $j$ is here defined as the eigenvector of $\mathbf{Y}_j^T\mathbf{Y}_j$ corresponding to the largest eigenvalue. While not strictly necessary for the $k$-means algorithm described above, it is typical to normalize the resulting centroids to unit vector norm (Pascual-Marqui et al., 1995), a process which is performed automatically in many implementations of eigenvalue decomposition, including the implementation used here (Matlab's eig function).

With the updated cluster centroids, the algorithm returns to step #2 (Main Text Table 2), iteratively recalculating distances between maps and centroids, relabelling states, and recalculating the centroids. These steps are iterated until convergence is met, i.e. once there is no change in assigned cluster labels between subsequent iterations (Koenig et al., 1999), or up to a fixed maximum number of iterations, here chosen to be 100 iterations (Tait et al., 2020). Since the choice of initial maps is random, different runs of the algorithm may give different results on the same dataset. For this reason, we repeat the clustering 20 times using a different initial seed on each repeat (Koenig et al., 1999; Tait et al., 2020). The repetition with the highest globally explained variance (Murray et al., 2008, GEV) is used in subsequent analysis, where GEV is defined as:

$$\text{GEV} = \frac{\sum_i \sigma_i^2 \max_j(R^2(\mathbf{y}_i, \mathbf{c}_j))}{\sum_i \sigma_i^2}, \tag{6}$$

where $\sum_i$ denotes summation over samples, $\sigma_i$ is the GFP at sample $i$ (Equation 1), and $R(\mathbf{y}_i, \mathbf{c}_j)$ is the similarity between the map at sample $i$ and the centroid of cluster $j$ (Equation 2). It should be noted here that the phrase GEV is maintained in this study to align with microstate literature, but in reality this measure is only a true measure of explained variance (by a linear model) when $R^2$ is squared correlation, i.e. the case of sensor EEG signals. However, interpretation is highly similar for all modalities. GEV is always in the range $[0, 1]$, and when GEV=1, the cluster centroid maps can fully explain the data, while GEV=0 indicates the maps do not explain any data.

In the EEG microstate literature, a range of methods exist for clustering states (Michel and Koenig, 2018). The most widely used alternative to $k$-means is topographic atomize and agglomerate heirarchical clustering (TAAHC). TAAHC has been demonstrated to give consistent spatial topographies (Khanna et al., 2014) and information-theoretically invariant temporal sequences (von Wegner et al., 2018) when compared to $k$-means in EEG microstate sequences. TAAHC is notably more computationally expensive than $k$-means for large datasets (von Wegner et al., 2018), motivating our choice for $k$-means in this study. However, the TAAHC algorithm suffers from the same technical limitations when applied to source/MEG data as the original $k$-means formalism, and therefore the spatial transformations and generalized definitions of GFP and map (dis)similarity presented here may also be used to adapt the TAAHC algorithm for source/MEG microstates.

#### S1.2 Optimum number of microstates

A number of methods exist for choosing the optimum number of clusters/microstates from the data (Michel and Koenig, 2018). The most widely used in the EEG microstate literature are the Cross Validation Index (CVI) (Pascual-Marqui et al., 1995) and the Krzanowski-Lai (KL) criterion (Murray et al., 2008). CVI is highly dependent on number of sensors/ROIs, and is known to perform poorly for large numbers of ROIs ($\geq 64$ according to Murray et al. (2008)). Here, we used 274 channel MEG mapped to 230 source space ROIs, so we did not use CVI. While KL criterion has been used for resting-state microstate analysis (Hatz et al., 2015; Tait et al., 2020), it is believed to work appropriately for event-related potential analysis but result in spurious peaks for resting-state data Michel and Koenig (2018). Alternatively, Custo et al. (2017) considered eleven different criteria for the optimum number of microstates and developed a meta-criterion combining these solutions.

Here, we use the kneedle algorithm (Satopää et al., 2011) to choose the number of states. As the number of clusters $k$ increases, so does the variance explained. Increasing $k$ up to the true number of states will only give a large increase in GEV as more states of the brain are accounted for, whereas increasing $k$ above the true number of states only explains noise and hence will give marginal increases to GEV. Therefore the plot of GEV vs $k$ has a characteristic 'knee' shape. The kneedle algorithm aims to find the number of clusters in the dataset by finding the value of $k$ for which this knee occurs. This criterion is simple, intuitive, and easy to interpret. It does not depend on number of channels, and does not generate multiple spurious peaks due to the smooth increase of GEV with number of clusters. For this reason, we here used kneedle for the optimum number of states.

#### S1.3 Simulations

Simulations were performed to test the ability of the microstate methodology to estimate a ground truth microstate sequence and spatial maps. Simulations were repeated for $k = 4$ and $k = 10$ states. The three main steps to simulations are described below.

Firstly, a generative microstate sequence was simulated. While any sequence of numbers could be used here, we aimed to generate a sequence with EEG-microstate like properties and avoid assumptions known to be invalid for EEG microstates such as Markovianity or stationarity. To do so, we used a random walk decision tree approach, which reverse engineers the method commonly used to transform EEG microstate sequences to random walks using a decision tree (Van De Ville et al., 2010; von Wegner et al., 2016, 2018). $k - 1$ random walks were generated with Hurst exponent $0.7$ (Van De Ville et al., 2010). Each random walk was then converted to a binary sequence based on the sign of their derivative, i.e. +1 if they are increasing, and -1 for decreasing. Each binary sequence corresponds to a branch of a decision tree, which places a particular time point within a state. For the example of four states, the first split separates the states {1,2} from states {3,4}, while the second split separates states 1 and 2, and the third split separates states 3 and 4. For example, consider a time point with the three binary sequences having values of (+1,-1,+1). The first binary sequence is +1, so the current state is either state 1 or 2 (a value of -1 would indicate state 3 or 4). To decide which of these two states is the current sample, we consider the 2nd sequence (which separates states 1 and 2; if the first sequence had a value of -1 we could consider the 3rd sequence which separates states 3 and 4). Values of +1/-1 in the 2nd sequence correspond to states 1 and 2 respectively, so since the 2nd sequence had a value of -1 the state at this time point is 2. Prior to binarizing the states, we smoothed the random walk with a moving average filter. The width of the filter was chosen for each simulation such that the mean duration was approximately 50 ms (Tait et al., 2020). We will denote the resulting sequence $L$, which is a vector of length $T$ (the number of samples), where the $t$'th sample $L(t) \in \{1, 2, \ldots, k\}$. This method resulted in generative sequences with mean duration of approximately 50 ms, Hurst exponents of approximately 0.7-0.8, and which were highly non-Markovian in line with past EEG microstate studies (Van De Ville et al., 2010; von Wegner et al., 2017, 2018; Tait et al., 2020).

Neural dynamics were then generated for each microstate. Each state was assigned a Wilson-Cowan neural mass model in a parameter regime previously used to model resting-state dynamics (Deco et al., 2009; Abeysuriya et al., 2018). For state $j$, the equations of the model were

$$\tau_E \dot{E}_j = -E_j + \phi(w_+ E_j - I_j + P_0 + P_{ms}(t)) + \sigma \nu_E(t) \quad (7)$$

$$\tau_I \dot{I}_j = -I_j + \phi(w_I E_j) + \sigma \nu_I(t) \quad (8)$$

$$P_{ms}(t) = \begin{cases} P_1 & \text{if } L(t) = j \\ 0 & \text{otherwise} \end{cases} \quad (9)$$

Here, $E_j$ and $I_j$ correspond to the firing rates of pools of excitatory and inhibitory neurons associated with microstate $j$ respectively. $\nu_E$ and $\nu_I$ are independent Gaussian noise terms with variance 1. The sigmoid function $\phi(x)$ is given by

$$\phi(x) = \frac{c}{1 + \exp(-a(x - b))}. \quad (10)$$

The remaining parameters and their interpretations are given in Table S2.

The input $P_0$ was chosen by Deco et al. (2009) such that (in the absence of microstate related drive $P_{ms}$) the model has a single stable fixed point close to a Hopf bifurcation, and as such exhibits low amplitude aperiodic noise-driven fluctuations with resonance at an alpha band frequency. Therefore all $k - 1$ neural masses with $P_{ms} = 0$ at any given time are in this regime. When microstate associated drive is added ($P_{ms} = P_1$), the system moves across the Hopf bifurcation into a high-amplitude stable periodic alpha oscillation. The LFP of each microstate was given by the sum of synaptic input onto the excitatory population,

$$s_j(t) = w_+ E_j(t) - I_j(t), \quad (11)$$

where $s_j$ is the observed LFP for microstate $j$.

Simulations were performed using the stochastic Heun algorithm with time step of 0.1ms. Each simulation was initialized with 2 seconds to reach steady states, and then 60 seconds of data was simulated. Simulated

| Parameter | Description | Value |
|---|---|---|
| $\tau_E$ | Excitatory time constant | 10 ms |
| $\tau_I$ | Inhibitory time constant | 20 ms |
| $w_+$ | Strength of recurrent excitation | 1.4 |
| $w_I$ | Strength of excitation onto inhibitory neurons | 1.5 |
| $P_0$ | Background input to excitatory neurons | 10 |
| $P_1$ | Microstate associated input to excitatory neurons | 3 |
| $\sigma$ | Standard deviation of neural noise | 0.1 |
| $c$ | Maximal firing rate | 100 |
| $b$ | Firing threshold | 40 |
| $a$ | Slope of sigmoid | 0.1 |

Supplementary Table S2: Parameters of the neural mass model

data was subsequently downsampled to 256 Hz in line with the empirical data.

Finally, the states were assigned spatial maps. As with the choice of generative sequence, the choice of spatial map is arbitrary. Here we used open acess fMRI derived resting-state network maps (Smith et al., 2009) aligned to our 230 ROI atlas by averaging all voxels within an ROI. The result was a matrix $W \in \mathbb{R}^{230\times k}$, whose columns are the spatial maps for each state. If we define $S \in \mathbb{R}^{k\times T}$ to be a matrix containing the neural dynamics of each state, i.e. $S_{j,t} = s_j(t)$, then the simulated LFP at the 230 ROIs were given by $X = WS + \lambda\eta$, where $X \in \mathbb{R}^{230\times T}$ has rows corresponding to time series for each ROI. $\eta \in \mathbb{R}^{230\times T}$ is pink noise, and $\lambda$ is a scalar factor for the noise chosen such that SNR was 1.

##### S1.3.1 Comparison with other state-estimation methodologies

Because simulations have ground truth maps and microstate sequences, they were used to compare $k$-means against other clustering algorithms such as PCA, ICA (using the fastica algorithm (Hyvarinen, 1999)), and hidden Markov modelling (HMM, using the HMM-MAR toolbox `https://github.com/OHBA-analysis/HMM-MAR`). We also compared whether results were more appropriate when performed on raw source data or alpha band filtered amplitude envelopes. For $k$-means, this meant using the 'Source' or 'Amplitude' transformations (Table S1), while for HMM we used the recommended settings for raw source and amplitude data in the HMM toolbox, which are derived from (Baker et al., 2014) for amplitude data and (Vidaurre et al., 2018) for raw source data. Four statistics were used to assess the quality of the state estimation. Firstly, GEV was used as a purely data driven quantity. Secondly, we correlated the estimated spatial maps for each state against the ground truth maps using a template matching algorithm. Third, we assessed the temporal sequence accuracy using the normalized mutual information (MI) between the ground truth and estimated sequence. Finally, we also calculated the normalized mutual information only at the GFP peaks (MIg), since these are the samples with locally maximal SNR.

#### S1.4 Microstate segmented functional connectivity

Our method is described below, and a visual overview of the pipeline is given in Figure S1. We first assign each time point in the broadband (1-30 Hz) data a microstate class using the methods described above. We subsequently filter the data into one of three narrow bands (theta 4-8 Hz, alpha 8-13 Hz, beta 13-30 Hz), and calculate the analytical signal using the Hilbert transform. For a given microstate, we concatenate all samples of the analytic signal within this microstate class, and use this to derive time courses of instantaneous phases and amplitudes. While these time series are no longer continuous, the Hilbert transform was performed on the continuous data so they represent instantaneous phases and amplitudes of the continuous signal at these samples. Under the assumption that microstates are associated with specific patterns of phase synchronization (i.e. the hypothesis we wish to test), we can subsequently calculate functional connectivity from the concatenated phase/amplitude time courses. A large number of metrics exist to quantify functional connectivity. Since amplitude coupling is typically performed on a slower time scale through lowpass filtering and downsampling, here we chose to address microstate synchronization using phase coupling, specifically the weighted phase lag index (wPLI) (Vinck et al., 2011; Colclough et al., 2016), which is a reliable estimator of phase coupling (Colclough et al., 2016) while attenuating spurious connections due to leakage Vinck et al. (2011). The (unweighted) phase lag index was used by Hatz et al. (2015) in their study of microstate connectivity, but wPLI is more reliable and less biased due to small noisy perturbations than PLI (Vinck et al., 2011; Colclough et al., 2016). This method is repeated for each microstate class, to obtain functional connectivity patterns for each class.

The methodology described above assumes that microstates are associated with a unique pattern of phase synchronization, and therefore by concatenating samples within a particular class we obtain a stationary time course from which it is meaningful to calculate a single functional network. To test this hypothesis, we applied multi-variate pattern analysis (MVPA) using the MVPA-Light toolbox (Treder, 2020). For each frequency band, the following analysis was performed. Firstly, for each participant, scan, and microstate class, we followed the methodology above for calculating microstate functional connectivity. However, prior to calculation of wPLI, the concatenated phase/amplitude time-courses were segmented into non-overlapping windows of 1280 samples (equivalent to 5-seconds). Across all participants, scans, and microstate classes this resulted in 5413 networks per frequency. Each network was associated with a single microstate class. For each network, the weighted degree distribution (i.e. sum of all edges connecting to a node) was calculated, resulting in a 230 (number of ROIs) dimensional vector for each network. These 230 dimensional vectors were used as features in a multi-class linear discriminant analysis classifier, with the associated microstate class as the class-labels to be predicted. Classification used default values in the MVPA-Light toolbox, while 5-fold cross-validated classification accuracy was used as the metric of performance. If high classification accuracy is achieved, this is strongly suggestive that microstates are associated with unique patterns of phase synchronization, since it suggests that given any 1280 samples from a given microstate class (across participants/scans), this class can be predicted based on the functional connectivity pattern. Permutation testing was used to test for statistically significant classification accuracy, using 1000 permutations.

Finally, for visualization purposes only, we plotted the deviation of the microstate functional connectivity pattern from the static connectivity pattern. This was done since plotting raw microstate functional connectivity patterns gave visually similar results, suggesting an underlying static connectivity associated with all states. Firstly, we obtained a single microstate network (for each class) by averaging the network across all 5-second segments within the class, participants, and scans. We obtained a single static network (used for all classes) by segmenting the original data into 5-second epochs, calculating wPLI, and averaging the resulting networks over all epochs, participants, and scans. We subsequently normalized the microstate-connectivity by the static connectivity using the equation

$$\text{wPLI}^{ij}_{\text{deviation}} = \frac{\text{wPLI}^{ij}_{\text{microstate}} - \text{wPLI}^{ij}_{\text{static}}}{1 - \text{wPLI}^{ij}_{\text{static}}}, \tag{12}$$

where superscripts $i, j$ denote ROIs $i$ and $j$. This equation has in the past been used to remove linear-connectivity from non-linear connectivity matrices (Rummel et al., 2015), but here is applied to remove static connectivity from microstate-connectivity matrices. For visualization, the top 1% of edges from the matrix $\text{wPLI}_{\text{deviation}}$ are shown.

### S2 Supplementary Results

#### S2.1 Reproducibility and reliability of maps

To test the reliability and reproducibility of the estimated microstate maps, we repeated the $k$-means clustering 250 times using a different random seem and sampling a different selection of 150,000 GFP peaks. For each of these permutations, only a single repetition of the algorithm was performed (unlike the original run, where 20 repetitions were used and the repetition with highest GEV was chosen). The estimated 10 maps on each of these iterations were aligned to the original 10 maps using iterative template matching, i.e. map similarity between the estimated maps and the templates (the original 10 maps) was calculated according to Equation 2, and the template/map pair with highest similarity were given the same label. Then both this template and map were removed from the pool. This was repeated until all maps were aligned.

To quantify how tightly the 250 runs were clustered about the original maps, we use the Silhouette coefficient (Rousseeuw, 1987). The silhouette coefficient has a value for each sample (i.e. 250 repetitions and 10 clusters), ranging from -1 (poorly clustered) to +1 (well clustered). Distance between maps was calculated using Equation 4. The measure rewards low intra-cluster distances and high inter-cluster distances. Therefore

| Statistic | Value | $p$-stationary |
|---|---|---|
| GEV | $63.05 \pm 0.41\%$ | $3.1 \times 10^{-16}$ |
| MD | $59.86 \pm 1.09$ ms | $1.6 \times 10^{-7}$ |
| H | $0.6829 \pm 0.0103$ | $1.3 \times 10^{-3}$ |
| C | $0.5986 \pm 0.0134$ | $1.9 \times 10^{-11}$ |
| stdCov | $14.28 \pm 0.59\%$ | $0.5667$ |
| stdDur | $15.89 \pm 1.03$ ms | $5.8 \times 10^{-5}$ |
| stdOcc | $1.913 \pm 0.059$ s$^{-1}$ | $2.3 \times 10^{-7}$ |
| **Statistic** | **Value** | **Markov 95%** |
| $G_0$ | $2.881 \pm 0.091 \times 10^5$ | $103.0$ |
| $G_1$ | $1.242 \pm 0.049 \times 10^3$ | $877.3$ |

Supplementary Table S3: Statistics of MEG microstates

if the estimated maps for all 250 runs are highly repeatable, the template matching algorithm will place each run in approximately the same location in the abstract space spanned by the map values, resulting in low intra-cluster distances (and high inter-cluster distances relative to the low intra-cluster distances) and therefore large silhouette coefficient.

Results are shown in Figure S2. On average, there was high similarity between our microstate maps and maps estimated from subsequent runs, and the mean ($\pm$ standard error) of similarity values across all classes and runs was $0.9668 \pm 0.0019$. The silhouette coefficient was $0.7726 \pm 0.0047$, indicating that the various repetitions were well clustered. Of the maps, the frontal and left/right parietal were the least reproducible (still having silhouette coefficients $> 0.4$ and map similarities $> 0.83$, indicating high reliability).

#### S2.2 Assessing non-stationarity

An important point to note is that the $k$-means clustering algorithm will cluster the data even in the absence of true stable microstates. We therefore additionally tested whether the statistics of the microstate sequences could be replicated by random fluctuations in a stationary Gaussian process, or whether they required genuine non-stationarity and the appearance of stable states. To do so, for each participant and scan, we generated 100 multivariate iterative amplitude adjusted Fourier transform (mvIAAFT) surrogates (Schreiber and Schmitz, 2000). These surrogate time series are stationary Gaussian processes with the same power/cross spectra and distributions as the original dataset (Schreiber and Schmitz, 2000; Rummel et al., 2011). Therefore if the microstate statistics from the empirical MEG data were drawn from the same distribution as the microstate statistics from the surrogate data, it is probable that the observed 'microstates' are really just due to random fluctuations of a stationary process. Similar approaches have been taken to statistically analyse non-stationarity in dynamic functional connectivity in neuroimaging datasets (O'Neill et al., 2018). Here, we found no surrogate distributions were significantly different from a normal distribution (minimum $p$-value over all participants, scans, and microstate statistics was $p = 0.2310$ using a Kolmogorov-Smirnov test), so therefore quantified differences in the empirical statistics vs the surrogates via a $z$-score, i.e. $z = (x - \mu_{\text{surrogate}})/\sigma_{\text{surrogate}}$, where $x$ is the empirical statistic of interest, and $\mu_{\text{surrogate}}$ and $\sigma_{\text{surrogate}}$ are the mean and standard deviation of the surrogate distributions respectively. For each statistic, the $z$-scores were subsequently averaged over scans, giving a single $z$-value per participant. To test whether the resulting distribution of $z$-scores across participants was likely drawn from a normal distribution with mean zero (i.e. whether the $z$-scores were significantly different from the surrogates), a one sample $t$-test was used for each statistic. For this analysis, the statistics which were class-specific (duration, coverage, occurrence) were transformed to a standard deviation across classes, so the test estimated specifically whether heterogeneity between microstates in these properties could be accounted for in a stochastic process.

Results of the non-stationarity analysis are shown in Supplementary Table S3 and Supplementary Figure S3. Mean duration of microstates was significantly larger than could be explained by random fluctuations in a stationary Gaussian process with the same power spectra, cross-spectra (and covariances), and distribution of data, suggestive of non-stationarity and the occurence of stable microstates in empirical resting-state MEG. Heterogeneity in durations across states (quantified by standard deviation; stdDur) was also significantly larger in empirical data than due to chance.

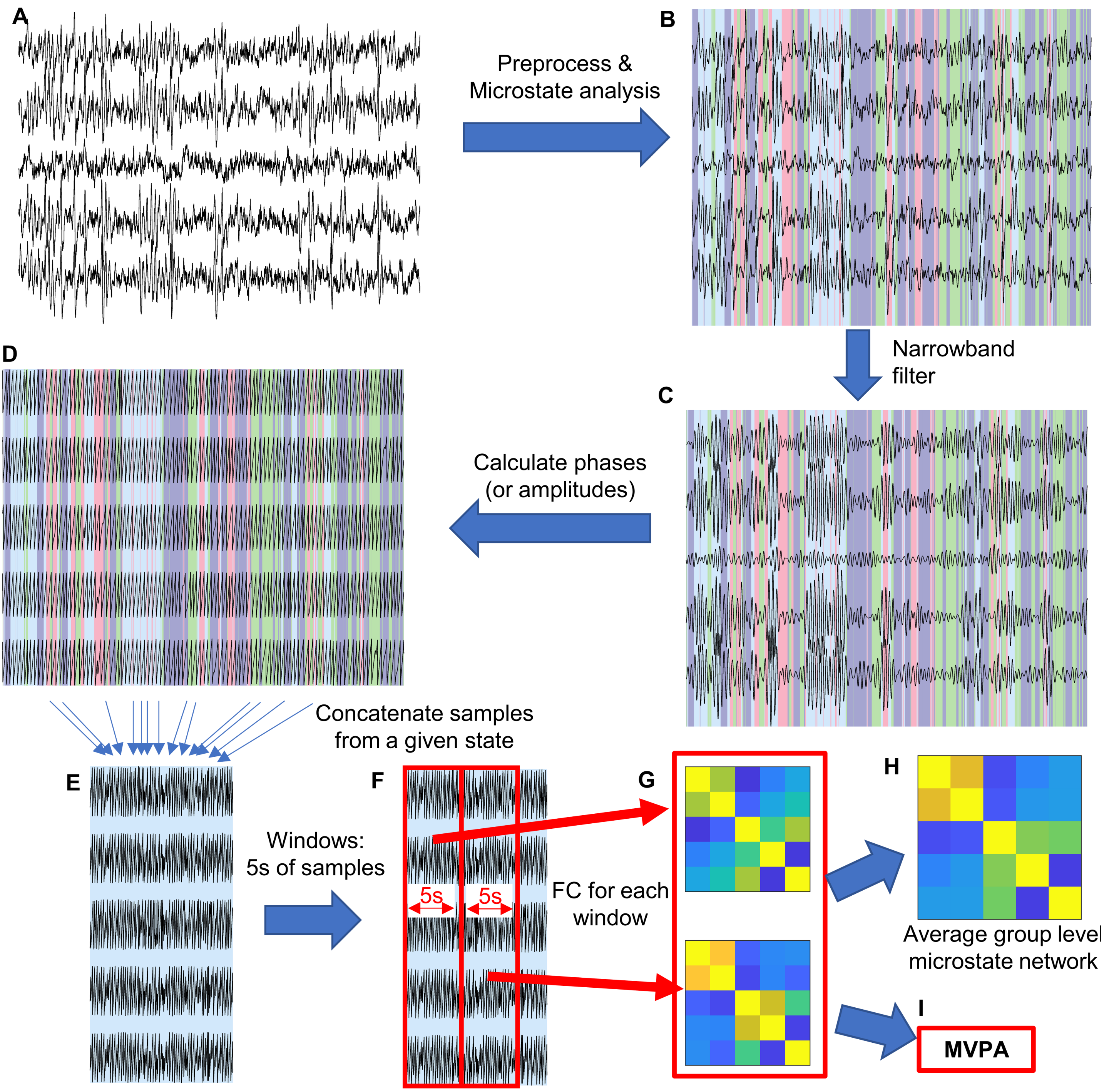


Supplementary Figure S1: **Calculating microstate functional connectivity**. (A) Raw MEG source space data. Note the data shown here is simulated data for visualization purposes, but empirical data was used in the analysis. (B) By appropriately preprocessing and performing microstate analysis on the data, we can assign each sample a microstate class (denoted here by colour). (C) To calculate narrowband functional connectivity, the data must be filtered into an appropriate frequency band. (D) Using the Hilbert transform, calculate instantaneous phases or amplitudes from the narrowband data. (E) Concatenate all samples with a label according to a given state. Here, we have shown all samples from the state with a light-blue background. (F) Window the concatenated data into epochs with 5 seconds worth of samples. (G) For each window, calculate functional connectivity. Here, we used wPLI. (H) To obtain a group level representation of the functional network associated with this microstate, average all functional connectivity matrices derived from this state. (I) To statistically test whether microstates are associated with a specific functional network structure, each matrix (along with the sets of matrices from all other microstate classes) can be submitted for multivariate pattern analysis.

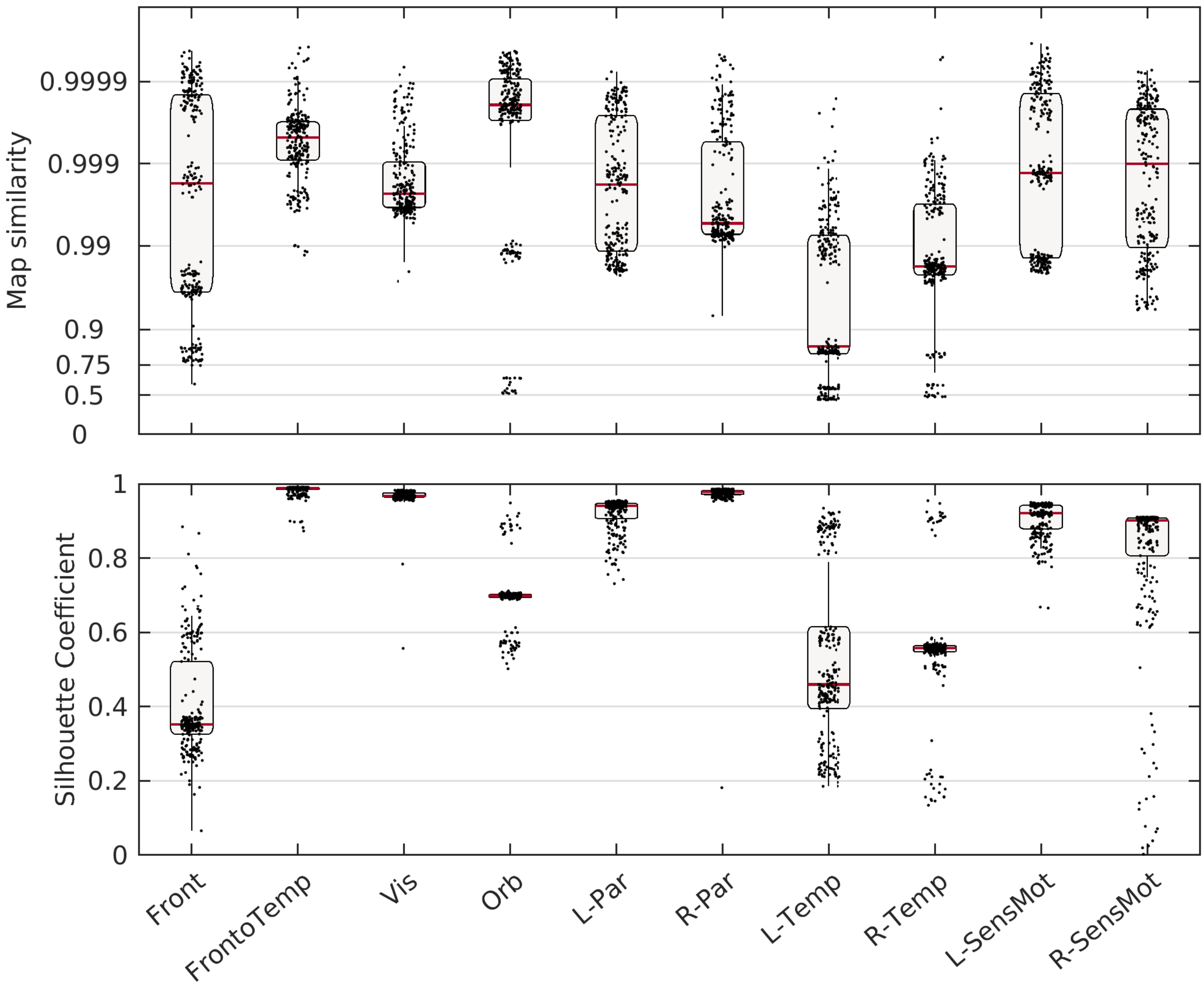


Supplementary Figure S2: **Reproducibility of resting-state microstates**. The $k$-means clustering algorithm was repeated 250 times with different sets of GFP peaks and initial seeds, and then matched to the original set of maps using a template matching algorithm. Each point correponds to a map estimated by a different repeat of the algorithm, while the classification according to the original maps is given by the x-axis. (Top) Map similarity between the estimated and original maps. The y-axis labels correspond to the raw values of map similarity, but the axis itself was transformed according to a Fisher $z$-transform to help differentiate large values of map similarity. (Bottom) Silhouette coefficients of each point.

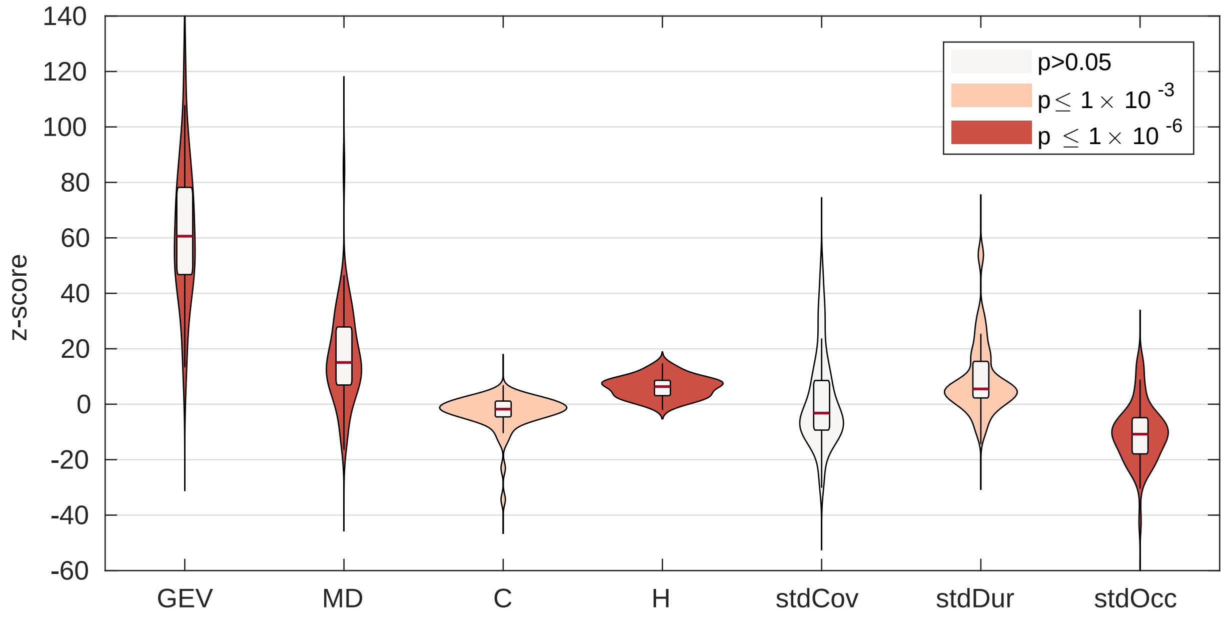


Supplementary Figure S3: **z-scores of empirical microstate statistics vs stationary surrogate distributions.**

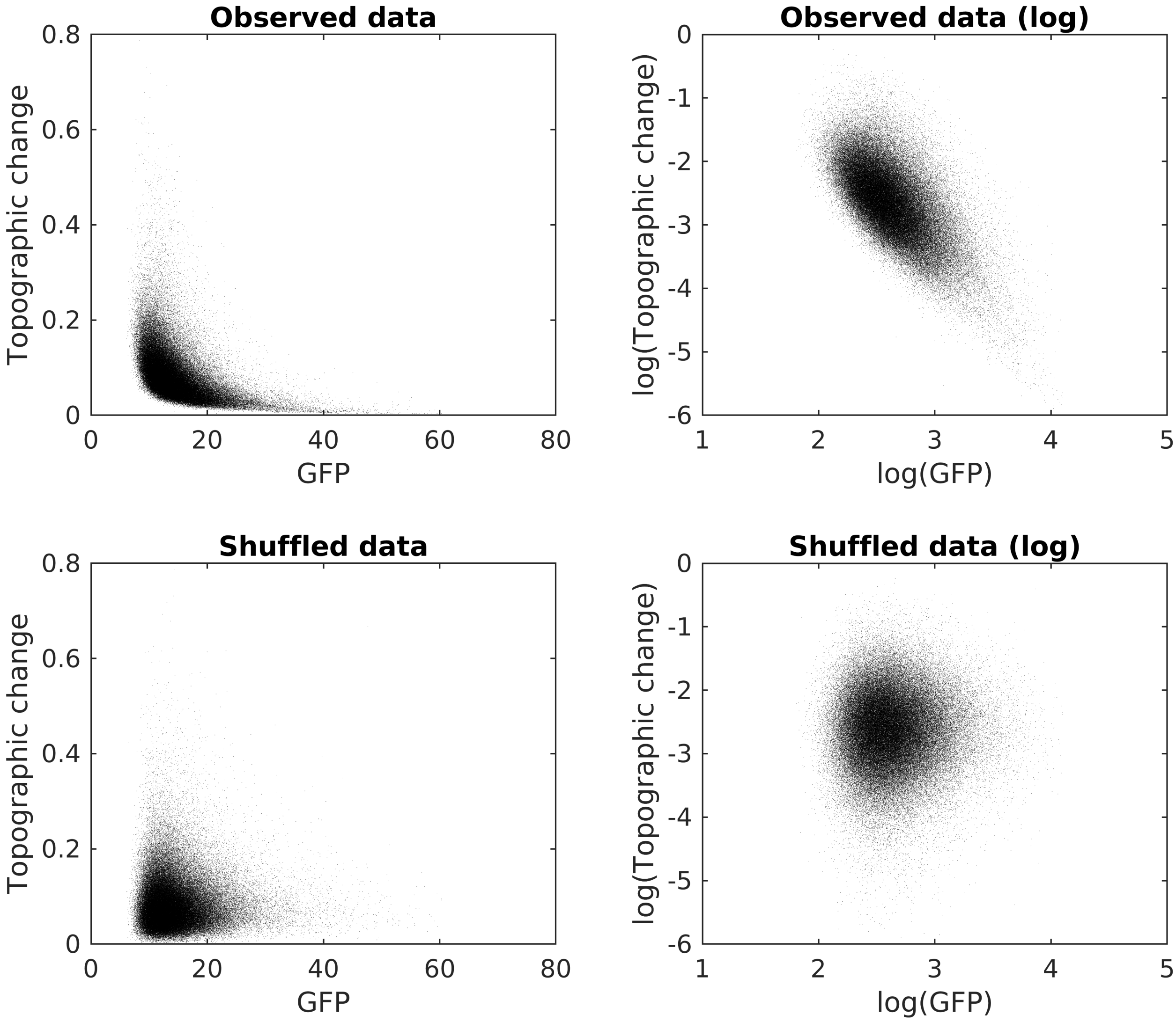


Supplementary Figure S4: **GFP vs topographic change.** Here, we recreate Fig 1 from Koenig and Brandeis (2016) using resting-state source MEG microstates, with modified definitions of GFP and map (dis)similarity as described in section S1.1. Upper rows show the observed data, while lower rows show shuffled data to simulate assumptions of Hidden Markov Modelling (i.e. independence of GFP and topographic change; Koenig and Brandeis (2016)). Left column shows original data, while right column shows log-log transformed data. After log-log transformation, there is a correlation of $-0.6225 \pm 0.0189$ (mean $\pm$ standard error across participants). The data shown here is from a single representative participant, with correlation of $-0.6466$. This figure demonstrates that the inverse relationship between GFP and the topographic change holds in MEG source space data, justifying the use of GFP peaks (as the most stable samples) in the $k$-means clustering algorithm and demonstrating that the assumptions of Hidden Markov Modelling may be inappropriate for this data (Koenig and Brandeis, 2016).
